# Transposable elements and their KZFP controllers are drivers of transcriptional innovation in the developing human brain

**DOI:** 10.1101/2020.12.14.422620

**Authors:** Christopher J. Playfoot, Julien Duc, Shaoline Sheppard, Sagane Dind, Alexandre Coudray, Evarist Planet, Didier Trono

## Abstract

Transposable elements (TEs) constitute 50% of the human genome and many have been co-opted throughout human evolution due to gain of advantageous regulatory functions controlling gene expression networks. Several lines of evidence suggest these networks can be fine-tuned by the largest family of TE controllers, the KRAB-containing zinc finger proteins (KZFPs). One tissue permissive for TE transcriptional activation (termed ‘transposcription’) is the adult human brain, however comprehensive studies on the extent of this process and its potential contribution to human brain development are lacking.

In order to elucidate the spatiotemporal transposcriptome of the developing human brain, we have analysed two independent RNA-seq datasets encompassing 16 distinct brain regions from eight weeks post-conception into adulthood. We reveal an anti-correlated, KZFP:TE transcriptional profile defining the late prenatal to early postnatal transition, and the spatiotemporal and cell type specific activation of TE-derived alternative promoters driving the expression of neurogenesis-associated genes. We also demonstrate experimentally that a co-opted antisense L2 element drives temporal protein re-localisation away from the endoplasmic reticulum, suggestive of novel TE dependent protein function in primate evolution. This work highlights the widespread dynamic nature of the spatiotemporal KZFP:TE transcriptome and its potential importance throughout neurotypical human brain development.

## Introduction

KZFPs constitute the largest family of transcription factors encoded by mammalian genomes. These proteins harbor an N-terminal Krüppel-associated box (KRAB) domain and a C-terminal zinc finger array, which, for many, mediates sequence-specific DNA recognition. The KRAB domain of a majority of KZFPs recruits the transcriptional co-repressor KAP1 (KRAB-associated protein 1, also known as Tripartite motif protein 28, TRIM28), which acts as a scaffold for heterochromatin inducers such as the histone methyl-transferase SETDB1, the histone deacetylating NuRD complex, heterochromatin protein 1 (HP1) and DNA methyltransferases (Ecco et al. 2017). Many KZFPs bind to and repress TEs, a finding that led to the ‘arms race’ hypothesis, which states that waves of genomic invasion by TEs throughout evolution drove the selection of KZFP genes after they first emerged in the last common ancestor of tetrapods, lung fish and coelacanth some 420 million years ago (Jacobs et al. 2014; Imbeault et al. 2017). While partly supportive of this proposal, functional and phylogenetic studies point to a more complex model, strongly suggesting that KZFPs have facilitated the co-option of TE-embedded regulatory sequences (TEeRS) into transcriptional networks throughout tetrapod evolution (Najafabadi et al. 2015; Imbeault et al. 2017; Helleboid et al. 2019). TEeRS indeed host an abundance of transcription factor (TF) binding sites (Bourque et al. 2008; Sundaram et al. 2014), and KZFPs and their TE targets influence a broad array of biological processes from early embryogenesis to adult life, conferring a high degree of species specificity (Trono 2015; Pontis et al. 2019; Chuong et al. 2013, 2016; Turelli et al. 2020). TEeRS can act as enhancers, repressors, promoters, terminators, insulators or via post-transcriptional mechanism (Garcia-Perez et al. 2016; Chuong et al. 2017). While these co-opted TE functions are key to human biology, their deregulation can also contribute to pathologies such as cancer and neurodegenerative diseases (Jang et al. 2019; Attig et al. 2019; Chuong et al. 2016; Li et al. 2015; Ito et al. 2020; Jönsson et al. 2020).

KZFPs and TEs are broadly expressed during human early development, playing key roles in embryonic genome activation and controlling transcription in pluripotent stem cells (Theunissen et al. 2016; Pontis et al. 2019; Turelli et al. 2020). However, how much TEeRS and their polydactyl controllers influence later developmental stages and the physiology of adult tissues is still poorly defined. Intriguingly, KZFPs are collectively more highly expressed in the human brain than in other adult tissues, suggesting a prominent impact for these epigenetic regulators and their TEeRS targets in the function of this organ (Nowick et al. 2009; Imbeault et al. 2017; Farmiloe et al. 2020; Turelli et al. 2020). In line with this hypothesis, we recently described how ZNF417 and ZNF587, two primate specific KZFPs repressing HERVK (human endogenous retrovirus K) and SVA (SINE-VNTR-Alu) integrants in human embryonic stem cells (hESC), are expressed in specific regions of the human developing and adult brain (Turelli et al. 2020). Through the control of TEeRS, these KZFPs influence the differentiation and neurotransmission profile of neurons and prevent the induction of neurotoxic retroviral proteins and an interferon-like response (Turelli et al. 2020). Furthermore, expression of LINE1, another class of TEs, has been noted in human neural progenitor cells (hNPCs) and in the adult human brain, occasionally leading to *de novo* retrotransposition events (Muotri et al. 2005; Coufal et al. 2009; Muotri et al. 2010; Upton et al. 2015; Erwin et al. 2016; Guffanti et al. 2018). Finally, various patterns of TE de-repression have been reported in several neurodevelopmental and neurodegenerative disorders, indicating that a de-regulated ‘transposcriptome’ may be detrimental to brain development or homeostasis (Tam et al. 2019; Jönsson et al. 2020).

A growing number of genomic studies relying on bulk RNA sequencing (RNA-seq), single cell RNA sequencing (scRNA-seq), assay for transposase accessible chromatin using sequencing (ATAC-seq) and other types of epigenomic analyses are teasing apart the transcriptional landscape of the developing human brain, revealing its dynamism and the complexity of the underlying cellular make-up (Kang et al. 2011; Miller et al. 2014; Fullard et al. 2018; Li et al. 2018; Keil et al. 2018; Zhong et al. 2018; Cardoso-Moreira et al. 2019). The present work was undertaken to explore the contribution of TEs and their KZFP controllers to this process. Our results identify KZFPs and TEeRS as important spatiotemporal contributors to gene expression in both the developing and adult brain, and reveal how neurological proteins with modified characteristics can arise from TE-mediated transcriptional innovations.

## Results

### Spatiotemporal patterns of KZFP gene expression during brain development

In order to determine the spatiotemporal patterns of KZFPs and TE expression in human neurogenesis, we analysed RNA-seq data from 507 samples corresponding to 16 different brain regions and 12 developmental stages (from 4 weeks post-conception to adulthood) available through the Brainspan Atlas of the Human Brain (Miller et al. 2014) and through Cardoso-Moreira et al. 2019 (Supplemental Fig. S1A & B). While the latter dataset comprises 114 samples exclusively from dorsolateral frontal cortex (DFC) and cerebellum (CB), transcriptomes for these regions were largely concordant with those documented in Brainspan, justifying the two resources as suitable for reciprocal validation (Supplemental Fig. S1C & D; Supplemental Table 1 & 2). We first examined KZFP gene expression in these two brain regions, which are representative of the forebrain and the hindbrain, respectively (Supplemental Fig. S1B). The large majority of KZFPs expressed in the DFC exhibited higher levels at early prenatal stages to drop shortly before birth and remain low onwards (Fig. 1A). When comparing early prenatal (2A-3B; 8-18 post-conception weeks) and adult (11; age 20-60+ years) stages, about half (169/333) of KZFPs were more expressed in the former and only 1.5% (5/333) in the latter, the rest being stable (Fig. 1B). This temporal pattern was less striking in the cerebellum (Supplemental Fig. S2A), with only 15.9% (53/333) and 2.1% (7/333) of KZFPs more strongly expressed in early prenatal and in adult respectively (Supplemental Fig. S2B). Thus, KZFP gene expression patterns are characterized by both temporal and regional specificity.

**Figure 1.**
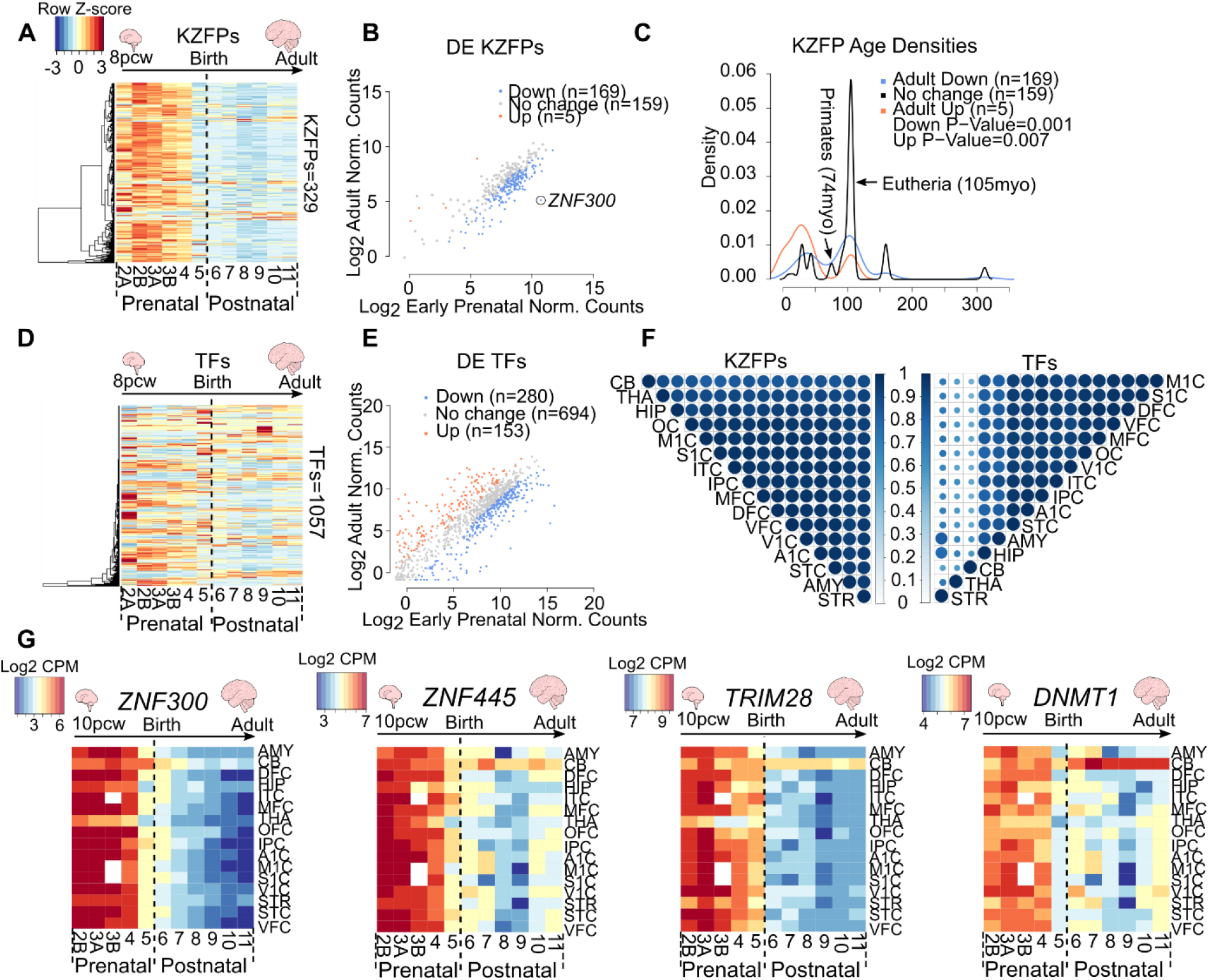
KZFP genes exhibit a global pre to postnatal decrease in expression. (A) Heatmaps of KZFP expression across human neurogenesis in the DFC. Scale represents the row Z-score. See also Supplemental Table 2 (B) Dot plot of differential expression analysis of KZFP genes in the DFC comparing adult (stage 11) to early prenatal stages (stage 2A to 3B) of neurogenesis. Only KZFPs differentially expressed in both datasets are shown. Up (orange) represents KZFPs significantly upregulated in adult versus early prenatal (Fold change ≥ 2, FDR ≤ 0.05). Down (blue) represents KZFPs significantly downregulated in adult (Fold change ≤ −2, FDR ≤ 0.05). See also Supplemental Table 3. (C) Density plot depicting estimated age of KZFPs of each category in (B) (*P*≤0.05, Wilcoxon test). (D) Heatmaps of TF expression across human neurogenesis in the DFC. Scale same as in A. (E) Dot plot of differential expression analysis of TFs (as defined in Lambert et al., 2018) in the DFC, excluding KZFP genes, comparing adult (stage 11) to early prenatal stages (stage 2A to 3B) of neurogenesis. Only TFs differentially expressed in both datasets are shown. Up (orange) represents TFs significantly upregulated in adult versus early prenatal (Fold change ≥ 2, FDR ≤ 0.05). Down (blue) represents KZFPs significantly downregulated in adult (Fold change ≤ −2, FDR ≤ 0.05). See also Supplemental Table 3. (F) Correlation plots representing the Pearson correlation coefficient of temporal KZFP expression (left) and TF expression (right) between all 16 regions. Size of spot and colour both represent the correlation coefficient. 0=no correlation, 1=strong correlation. (G) Heatmaps depicting the log2 counts per million (CPM) for selected KZFPs and TFs over the 16 regions included. See also Supplemental Table 1 & 2. All plots show expression data from Brainspan.

KZFP genes have emerged continuously during higher vertebrate evolution, collectively undergoing a high turnover in individual lineages. Amongst some 360 human KZFPs, about half are primate-restricted, whereas a few are highly conserved, with orthologous sequences present in species that diverged more than 300 million years ago (Imbeault et al. 2017; Huntley et al. 2006). To determine if the differentially expressed KZFPs arose at particular times in evolution, we determined their ages. We found KZFPs either significantly downregulated or upregulated from early prenatal to adult stages to be significantly younger than those displaying no differences between these developmental periods (Fig. 1C, Wilcox test *p*<=0.01). This delineates two subsets amongst KZFPs participating in brain development, one evolutionarily recent and more transcriptionally dynamic, the other more conserved and transcriptionally static.

Of note, KZFPs appeared distinct amongst TFs (as defined in Lambert et al. 2018), as other members of this functional family exhibited far more diverse patterns of expression throughout development, whether in the forebrain or in the cerebellum (Fig. 1D & E; Supplemental Fig. S2C & D). Only about a quarter of TFs were indeed more highly expressed in early prenatal stages in either region, against around 10% in the adult brain (Fig. 1E; Supplemental Fig. S2C & D). Furthermore, temporal expression patterns of KZFP genes were highly correlated across all 16 brain regions, albeit to a lesser extent in the cerebellum (Fig. 1F). In contrast, other TFs displayed far more diverse behaviours, with the CB, mediodorsal nucleus of the thalamus (THA) and striatum (STR) exhibiting reduced correlation values compared to other regions (Fig. 1F). Thus, KZFPs are collectively subjected to a remarkable degree of spatiotemporal coordination in spite of the diversity of their genomic targets and of cell types present in the various regions of the brain. The KZFP gene most differentially expressed in prenatal versus postnatal DFC was the hematopoietic differentiation associated *ZNF300* (Xu et al. 2010) (Fig. 1B; Supplemental Table 3). This was true in all brain regions, although its transcripts persisted longer in the cerebellum compared to other areas (Fig. 1G; Supplemental Table 1 & 2). *ZNF445*, which binds and controls imprinted loci in humans (Takahashi et al. 2019), similarly exhibited comparable patterns across all brain regions but its expression was largely maintained in the cerebellum all the way to adulthood (Fig. 1G; Supplemental Table 1 & 2).

We next examined *KAP1,* which encodes a protein that serves as corepressor for many KZFP (Ecco et al. 2017). Its expression levels were globally higher than those of any KZFP, albeit also with a drop from prenatal to postnatal stages except in the cerebellum (Fig. 1G; Supplemental Table 1 & 2). We also probed *DNMT1*, which encodes the maintenance DNA methyltransferase important for TE repression in neural progenitor cells and other somatic tissues beyond the early embryonic period (Jönsson et al. 2019). Although displaying overall patterns comparable to those seen for *KZFPs* and *KAP1*, *DNMT1* expression progressively increased in the cerebellum to reach its highest level in the adult (Fig. 1G; Supplemental Table 1 & 2). In sum, KZFPs and their main epigenetic cofactors exhibit a largely homogenous, dynamic spatiotemporal reduction in expression during human brain development.

### TE subfamilies are dynamically expressed throughout development

Having determined that the expression of most KZFPs drops at late stages of prenatal brain development, we examined the behaviour of their TE targets. Young TEs are highly repetitive, which complicates the mapping of TE-derived RNA-seq reads to unique genomic loci, thus biasing against the scoring of their expression. We therefore first analysed RNA-seq reads mapping to multiple TE loci within the same subfamily, regardless of positional information. In the DFC, discrete subfamilies, predominantly from the LTR class and to a lesser extent the SINE class, exhibited temporally distinct dynamics, concordant between datasets (Pearson correlation coefficient ≥ 0.7) (Fig. 2A; Supplemental Table 4). The same was true for the cerebellum, but with moderately different subfamilies passing our threshold for concordance between datasets (Supplemental Fig. S3A; Supplemental Table 4). In the DFC, for example, the LTR7C and SVA-D subfamilies exhibited higher postnatal expression, whereas LTR70 and HERVK13-int behaved inversely, albeit without marked differences between brain regions (Fig. 2B; Supplemental Table 4 & 5). Similarly to KZFP genes, TEs have emerged continuously throughout evolution, with both young integrants and relics of ancient TEs reflective of different waves of genomic invasion. Using TE subfamily age estimates from DFAM (Hubley et al. 2016), we found that dynamically expressed TEs, concordant between both datasets, were significantly younger than non-concordantly expressed subfamilies in the DFC and cerebellum (Fig. 2C; Supplemental Fig. S3B).

**Figure 2.**
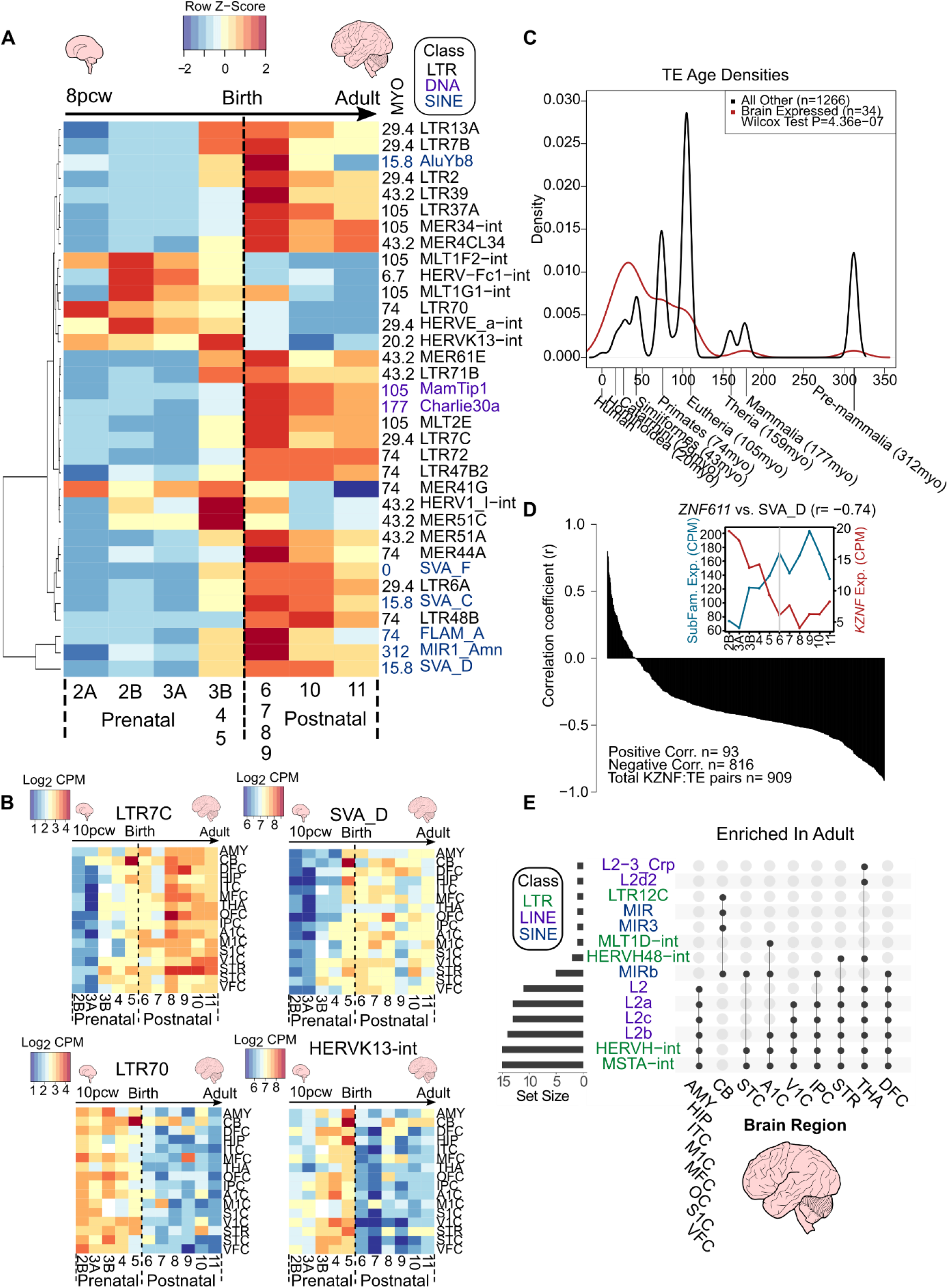
TE subfamilies and unique loci exhibit spatiotemporal expression patterns. (A) Heatmap of TE subfamilies with concordant expression behaviours between both datasets (Pearson correlation coefficient ≥ 0.7) across human neurogenesis in the DFC. See also Supplemental Table 4. The mean expression values for stages 3B, 4 and 5, and also stages 6, 7, 8 and 9 were combined and averaged to reduce inherent variability due to low numbers of samples for some stages (see Supplemental Fig. S1B). Scale represents the row Z-score. TE subfamily age in million years old (MYO) and class is shown to the right of the plot. (B) Heatmaps of TE subfamily expression across human neurogenesis in all 16 regions. See also Supplemental Table 4 & 5. Scale represents log2 CPM. Stage 2A was omitted due to lack of samples for some brain regions (see Supplemental Fig. S1B). (C) Density plot depicting estimated age of TEs in A (P≤0.05, Wilcoxon test). Evolutionary stages and corresponding ages are shown beneath the plot. (D) Barplot showing the Pearson correlation coefficient of KZFP expression and their target TE subfamily expression. 1=highly correlated, −1=highly anti-correlated. (D Inset) Line plot showing expression in counts per million of *ZNF611* and its main TE target subfamily, SVA_D and their Pearson correlation coefficient (−0.74, p-value=0.006). Grey line indicates birth at stage 6. See also Supplemental Table 6. (E) UpSet plot showing the significantly enriched differentially expressed subfamilies between adult and early pre-natal stages per region from unique mapping analyses. Set size represents the number of regions the specific TE was significantly differentially enriched in. Joined points represent combinations of significantly differentially expressed TE subfamilies. See also Supplemental Table 7 and 8. All plots show expression data from Brainspan.

We next analysed the temporal dynamics of the expression of KZFPs and their TE targets in the DFC, for which samples were available in highest abundance. For this, we matched KZFP ligands to their significantly bound TE subfamilies using an in-house algorithm on a large collection of ChIP-exo data (Imbeault et al. 2017). The results revealed that an overwhelming majority of KZFP:TE subfamily pairs (816 vs. 93) were anti-correlated in their expression, consistent with the known role of KZFPs as TE repressors (Fig. 2D; Supplemental Table 6). For example, *ZNF611* is a previously characterised major regulator of SVA-D in early embryogenesis (Pontis et al. 2019), and the two exhibited strongly anti-correlated expression throughout human brain development (Fig. 2D inset).

We next expanded our study by examining the expression of individual TE integrants, assigning RNA-seq reads to their genomic source loci and comparing early prenatal (stages 2A to 3B) and adult (stage 11) samples for the 16 available brain regions (Supplemental Fig. S1A & B). We found between 5,000 and 7,000 significant differentially expressed TE loci in each region, with 4,000 loci common to both DFC and CB datasets (Supplemental Fig. S3C; Supplemental Table 7 & 8). Integrants belonging to fourteen TE subfamilies from the LTR, LINE and SINE classes were significantly more expressed in adult samples, with HERVH-int, MSTA-int and L2 elements significantly enriched in most brain regions (Fig. 2E). The cerebellum again exhibited distinct patterns, with significant enrichment of LTR12C and MIR elements instead (Fig. 2E). Conversely, integrants from 11 TE subfamilies were more expressed in the early prenatal period, largely in specific brain regions (Supplemental Fig. S3D). Together, these results highlight the spatiotemporal dynamic nature of the transposcriptome in the developing human brain.

### Transpochimeric gene transcripts during human brain development

TE expression may be reflective of either ‘passive’ co-transcription from genic transcripts or *bona fide* TE promoter activity (reviewed in Lanciano and Cristofari 2020). Transpochimeric gene transcripts (TcGTs), that is, gene transcripts driven by TE-derived promoters, are the most easily interpretable and direct manifestation of the influence of TEeRS on gene expression. Some evidence for a role of TcGTs in the brain was provided by the recent observation that DNMT1 represses in hNPCs the expression of hominoid-restricted LINE1 elements, which subsequently act as alternative promoters for genes involved in neuronal functions (Jönsson et al. 2019). To explore more broadly the potential role of TcGTs in human brain development and function, we performed *de novo* transcript assembly, searching for mature transcripts with a TE-derived sequence at their 5’ end and the coding sequence of a cellular gene downstream. Due to the striking anti-correlation in KZFP and global TE expression between prenatal (stage 2A to stage 5) and postnatal stages (stage 6 to stage 11), we concentrated on these two periods, retaining only TcGTs present in greater than 20% of either prenatal, postnatal or both categories of samples and behaving in the same temporal manner in the two independent datasets. If there was a two-fold difference in the proportion of prenatal versus postnatal, the TcGT was annotated as either pre- or postnatal, whereas those below this threshold were deemed continual.

Our search yielded 480 high confidence TcGTs, of which 9.8% (47/480) were prenatal, 12.3% (59/480) postnatal and 72.3% (374/480) continual (Fig. 3A; Supplemental Table 9). Amongst pre- or postnatal TcGTs, developmental trajectories differed substantially, with some detected exclusively at either stage. For example, an L2a-driven isoform of *CTP synthase 2* (*CTPS2*), whose product catalyses CTP formation from UTP (van Kuilenburg et al. 2000), was found in 86% of all prenatal samples but only 12% of postnatal samples (Fig. 3A & B), whereas the inverse was observed for a MamGypLTR1b-driven isoform of the astrocyte associated *Aldehyde Dehydrogenase 1 Family Member A1* (*ALDH1A1*) (Adam et al. 2012) (12% vs. 95%) and an L2b-driven isoform of *Phospholysine Phosphohistidine Inorganic Pyrophosphate Phosphatase* (*LHPP*) (0.9% vs. 97%) (Fig. 3A), the host of intronic single nucleotide polymorphisms (SNPs) associated with major depressive disorder (Neff et al. 2009; Cui et al. 2016). The previously reported LTR12C-driven transcript of *Semaphorin 4D* (SEMA4D), the product of which participates in axon guidance (Cohen et al. 2009; Kumanogoh and Kikutani 2004), was detected in 79% of postnatal and only 0.9% of prenatal samples where it was instead expressed from a non-TE promoter, indicating a promoter switch during neurogenesis (Fig. 3A & B).

**Figure 3.**
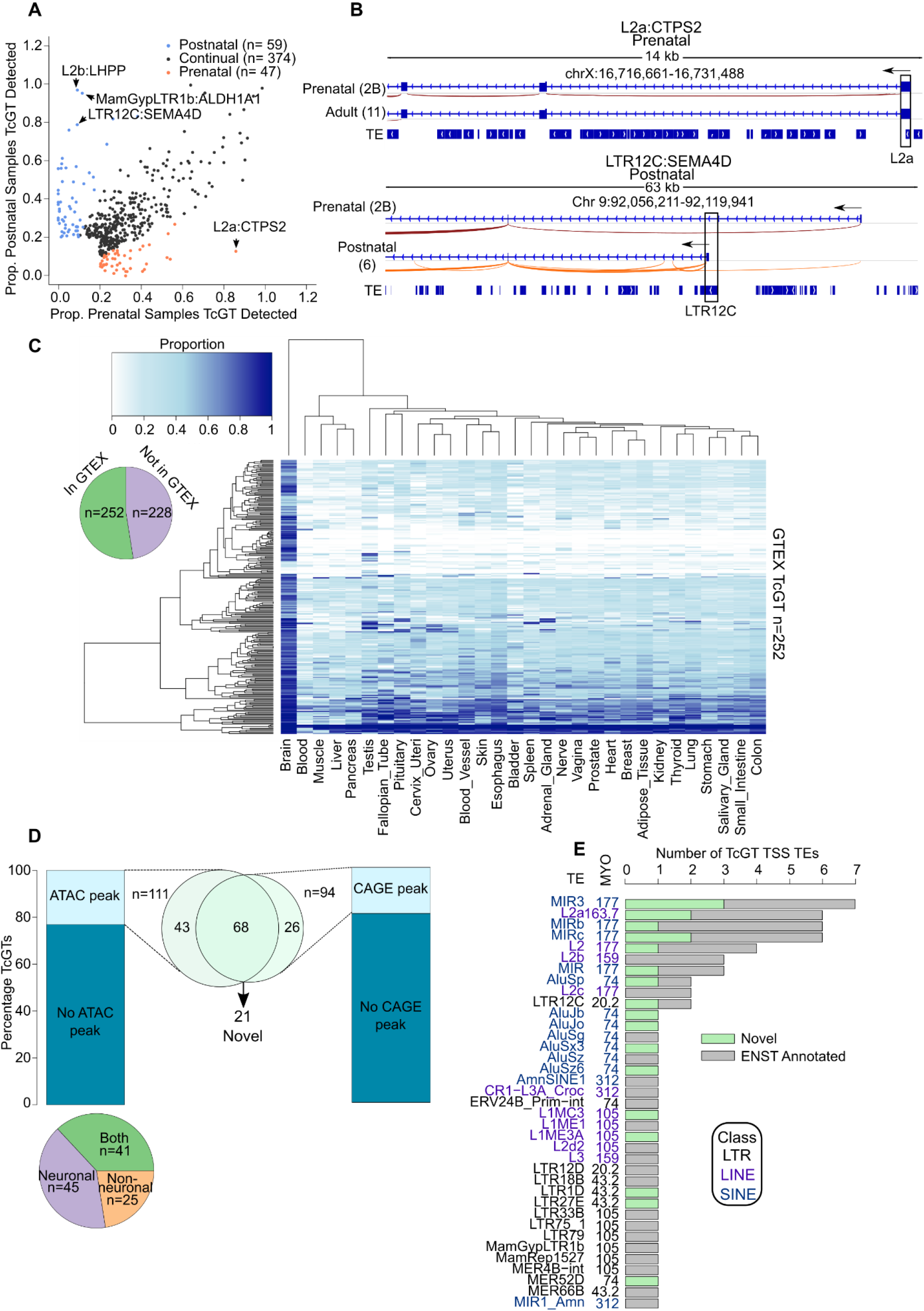
TE co-option as genic promoters drives spatiotemporal gene expression in human neurogenesis. (A) Dot plot showing the proportion of pre or postnatal samples TcGTs were detected in and behaving similarly in both datasets (prenatal, postnatal or continual). (B) Sashimi browser plots from IGV showing the splicing events in representative samples for prenatal enriched TcGT L2a:CTPS2 and the postnatal enriched LTR12C:SEMA4D. (C) Heatmap indicating the proportion of samples per GTEX tissue each TcGT from A was detected in. Each row represents an individual TcGT and each column a different tissue. (C inset) Pie chart indicating the proportion of neurodevelopmental TcGTs detected in GTEX. (D) Stacked barplot indicating the proportion of TcGT TE TSS loci overlapping an ATAC-seq peak from BOCA (left) and a pie chart indicating their cell type distribution (bottom left). Stacked barplot (right) indicating the proportion of TcGT TE TSS loci overlapping a FANTOM5 defined CAGE peak. Pie chart (centre) showing the overlap of ATAC-seq and CAGE peak associated TcGTs and highlighting 21 novel, non-ENSEMBL annotated transcripts. (E) Stacked barplots indicating the TE subfamily, TE class, TE age and the ENSEMBL overlap of each TcGT TE TSS loci. See also Supplemental Table 9 for all TcGT information.

We next examined the broader expression pattern of the 480 TcGTs detected during brain development. By applying our pipeline to the Genotype Tissue Expression (GTEX) dataset (Melé et al. 2015), we detected around half of them in this collection of predominantly adult samples (Fig. 3C; Supplemental Table 9). Some were present in all available tissues, but the vast majority were brain restricted (Fig. 3C).

### TcGTs exhibit cell type-specific modes of expression

We next analysed the state of the chromatin at the transcription start site (TSS) of the 480 TcGTs expressed during brain development by intersecting their proximal, TE-residing TSS (+/-200bp) with ATAC-seq consensus peaks from neuronal (NeuN+) and non-neuronal (NeuN-) cells across 14 distinct adult brain regions from the Brain Open Chromatin Atlas (BOCA) (Fullard et al. 2018). About a quarter (111/480) of these TcGTs TSS overlapped with ATAC-seq peaks in the adult brain, indicating that their chromatin was opened in this setting (Fig. 3D). Of these, two-thirds exhibited cell type specificity, either to neurons (40.5%, 45/111) or to non-neuronal cells (22.5% 25/111), whereas a third (41/111) were present in both cell subsets (Fig. 3D; Supplemental Table 9). These cell-restricted patterns were generally independent of the brain region considered, as illustrated by two postnatal enriched TcGTs, the non-neuronal L2-driven *Dysferlin* (*DYSF*) (Supplemental Fig. S4A), a gene mutations of which are associated with limb girdle muscular dystrophy 2B (Bashir et al. 1998; Liu et al. 1998), and the neuronal L2a-driven *Potassium Voltage-Gated Channel Subfamily A Regulatory Beta Subunit 2* (*KCNAB2*) encoding a regulator of neuronal excitability (McCormack et al. 2002) (Supplemental Fig. S4B).

To confirm that transcription of the TcGTs detected in the developing human brain was starting at the identified TE, we intersected their TSS with CAGE (cap analysis of gene expression) peaks previously defined in around 1,000 human cell lines and tissues by the FANTOM5 consortium (Forrest et al. 2014; Lizio et al. 2015). About a fifth of the TcGTs TSS (19.5%, 94/480) overlapped with CAGE peaks, of which 68 also corresponded to ATAC-seq peaks, providing a subset of high confidence TE-derived TSS loci driving gene transcription in the developing brain (Fig. 3D; Supplemental table 9). Of these, 21 were not annotated in ENSEMBL (Fig. 3D; Supplemental table 9), indicating that co-opted TEs acting as promoter elements are contributing to a previously undetected TE-derived neurodevelopmental transcription network.

We concentrated deeper analyses on the 68 high confidence brain developmental TcGTs. Thirty-seven different TE subfamilies accounted for their promoters but MIRs and L2s, belonging respectively to the SINE and LINE families, contributed almost half, perhaps due in part to their high prevalence in the genome (MiR3 and L2a: 87,870 and 166,340 integrants, respectively) (Fig. 3E), and LTRs about a fifth. A large range of evolutionary ages were represented, from the ~20 myo (million year old) LTR12C to the ~177 myo MIRs and L2s.

Of these 68 high-confidence TcGTs, 38.2% (26/68) were postnatal-specific, 51.5% (35/68) were continually detected and 10.3% (7/68) were prenatal-restricted (Fig. 4A). Furthermore, the 5’ end of these TcGTs coincided with ATAC-seq peaks from neurons in 26.5% (18/68), from non-neuronal cells in 22% (15/68), and from both in 51.5% (35/68) of cases (Fig. 4A). Some TcGTs were present in all brain regions, whereas others exhibited regional specificity (Supplemental Fig. S5A). For example, L2:DDRGK1 and L2a:KCNAB2, among others, were detected both postnatally and in a higher proportion of neocortex regions compared to the cerebellum (Fig. 4A; Supplemental Fig. S5A). We next aimed to determine if the detected TcGTs had the capacity to code for protein. Importantly, *in silico* prediction of the protein coding potential of these TcGTs, found that about half (31/68) likely encoded the canonical protein sequence and a fifth (15/68) an N-truncated isoform, while other configurations (N-terminal addition, C- or N- and C-truncation) were less frequent (Fig. 4A; Supplemental Table 9).

**Figure 4.**
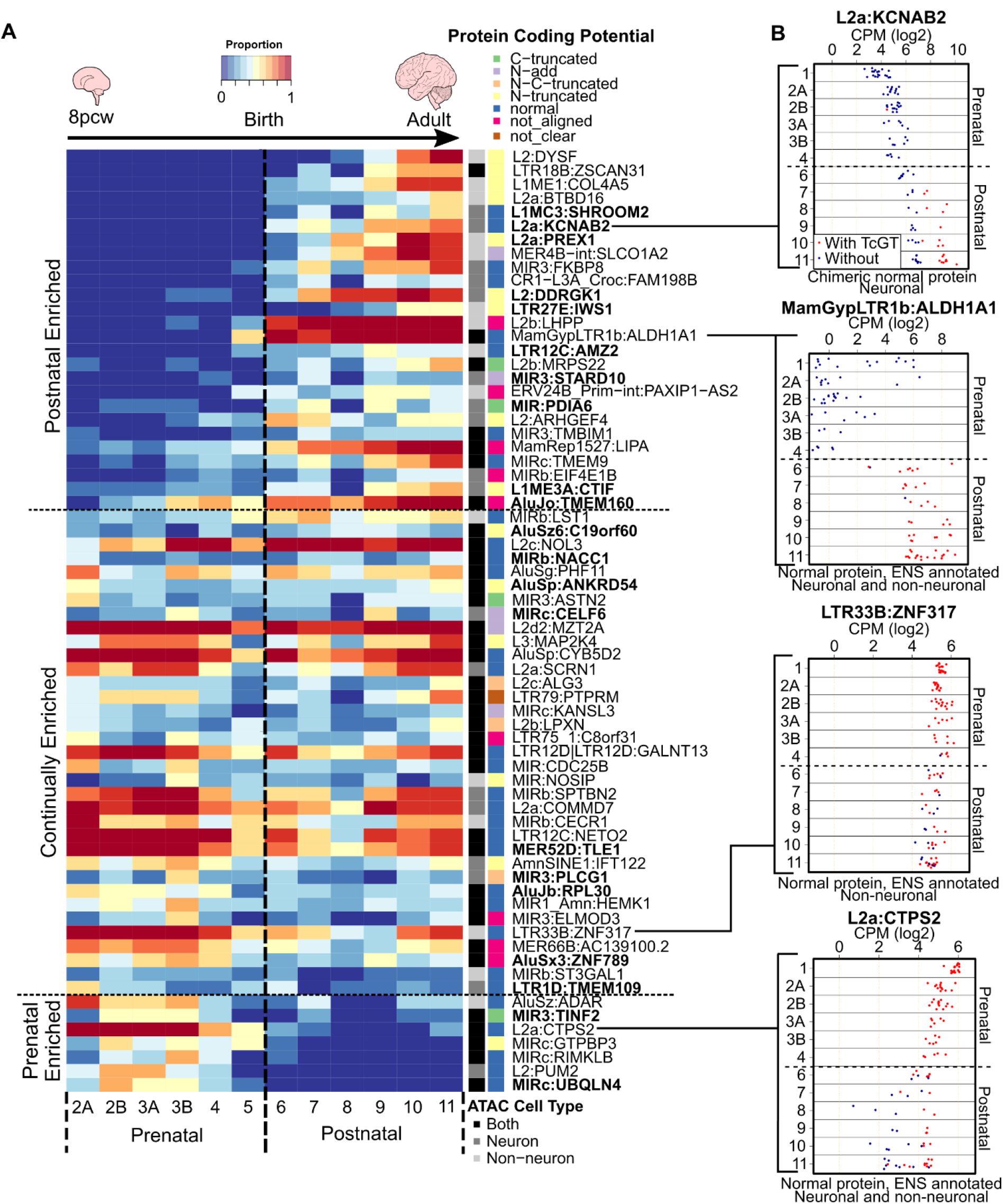
TcGTs are temporally expressed throughout neurogenesis in a cell type specific manner, exhibit protein coding potential and drive transcript expression. (A) Heatmap showing the proportion of samples per developmental stage the 68 TcGTs (from Fig. 3D) were detected in the Brainspan dataset, alongside their ATAC-seq cell type overlaps and protein coding potential determined via *in silico* translation. Bold indicates novel transcripts not annotated in ENSEMBL. See also Supplemental Table 9. (B) Dot plots showing the gene expression level per stage for the specified gene for samples where the TcGT was detected (red) and where it was not (blue) from Cardoso-Moreira dataset as comparison to (A). Dashed line represents birth at stage 6.

To estimate the relative contribution of the TE and non-TE promoters to the expression of the 68 genes involved in high confidence TcGTs, we compared their transcription levels in samples where the TcGT was or was not detected (Fig. 4B). In some cases, the TcGT was associated with higher levels of gene expression in a temporal manner such as the postnatally detected L2a:KCNAB2 (top) and most strikingly MamGypLTR1b:ALDH1A1 (top mid), compared to their non-TE-driven counterparts (Fig. 4B). The continually detected, non-neuronal LTR33B:ZNF317 (bottom mid) was associated with high expression throughout brain development, suggestive of a constitutive TE derived promoter. Conversely, some TcGTs were associated with higher prenatal expression, such as with L2a:CTPS2 (bottom), while for other genes there were more moderate expression differences in samples with and without TcGT detection as seen for the postnatally detected neuronal TcGT L2:DDRGK1 (Supplemental Fig. S5B).

### Experimental validation of brain-detected TcGTs

To verify that the TE and genic exon belonged to the same mRNA transcript, we next aimed to experimentally confirm TcGT candidates in the SH-SY-5Y neuroblastoma cell line. Using qRT-PCR primers within the TE TSS and subsequent genic exon, we detected appreciable expression of TcGTs in this cell system (Supplemental Fig. S6A). However, this did not formally demonstrate that transcription was driven by the TE. To address this point, we targeted a CRISPR-based activation system (CRISPRa) to the TSS region of TcGTs in 293T cells (Chavez et al. 2015) (Fig. 5A). We picked candidates based on the ease of gRNA design and the potential mechanistic or biological relevance of their protein product. We selected three anti-sense L2-driven, cell type-specific TcGTs predicted to encode for proteins involved in brain development: KCNAB2, DYSF and DDRGK1, the first in its canonical protein isoform and the other two as N-truncated isoforms. Activation of each of these three TcGTs could be induced with the CRISPRa system, confirming that they were indeed driven by their respective TE promoters (Fig. 5A).

**Figure 5.**
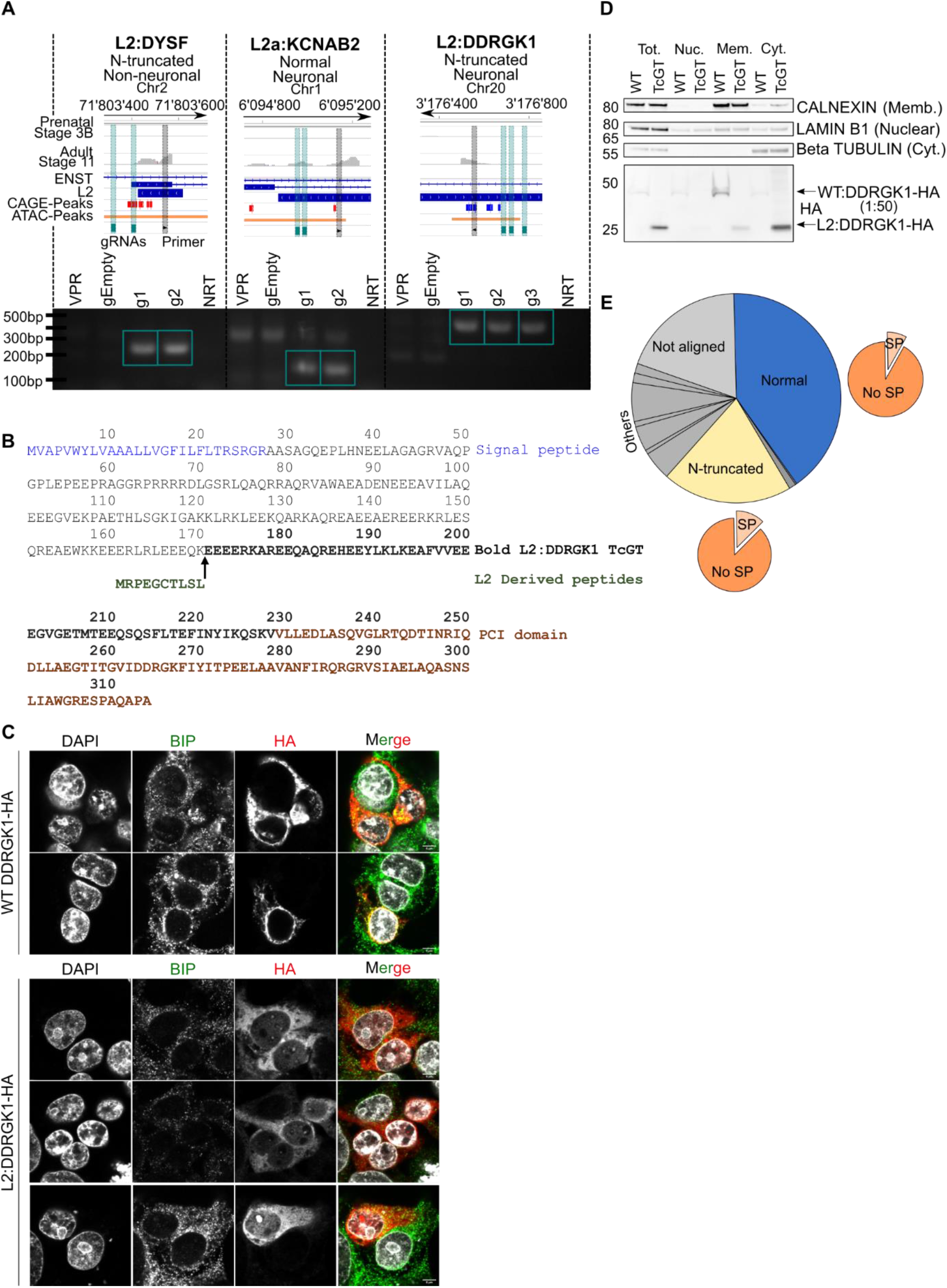
Antisense L2 elements directly drive TcGTs and contribute to chimeric protein formation and cytosolic re-localisation of the ER-membrane associated DDRGK1. (A) Schematic of TcGT TE TSS loci for indicated genes and representative prenatal (stage 3B) and adult (stage 11) RNA-seq tracks. Their associated protein coding potential and cell type specificity are highlighted and CAGE peak loci (red sense strand, blue anti-sense strand), CRISPRa gRNAs (green vertical bar) and TE associated PCR primers are shown (black vertical bar) (top). RT-PCR on cDNA generated from HEK293T cells transiently transfected with dCAS9-VPR plasmid and individual gRNA plasmids containing sequences targeting the TcGT TE TSS loci denoted in the schematic. dCAS9-VPR (VPR) or empty gRNA plasmids (gEmpty) alone were used as controls. Green box indicates bands of correct PCR product size absent in controls. NRT= no reverse transcriptase. (B) Canonical DDRGK1 and TcGT L2:DDRGK1 derived protein sequence. (C) Overexpression of canonical WT DDRGK1-HA and L2:DDRGK1-HA in HEK293T cells followed by immunofluorescent staining for BIP (an ER-membrane associated protein) and HA tag, followed by confocal imaging (scale bar = 5μm). (D) Overexpression of canonical DDRGK1-HA (WT) and L2:DDRGK1-HA (TcGT) in HEK293T cells followed by cellular fractionation and western blot for the indicated marker proteins (right of western blot) and HA tag. For WT DDRGK1 50x less protein lysate compared to L2:DDRGK1 was loaded for the HA blot due to high levels of protein expressed. Image is representative of two independent experiments. (E) Pie charts showing the *in silico* protein coding potential of the 480 TcGTs identified in Fig. 3A with the proportion containing a signal peptide shown with the orange pie charts. See also Supplemental Table 9.

### TcGT-encoded protein isoforms can display differential subcellular localisation

Having noted that 22% of high-confidence TcGTs were predicted to encode N-truncated proteins (Fig. 4A), we hypothesised that this could, in some cases, result in derivatives deprived of important subcellular localization domains, such as the endoplasmic reticulum (ER)-targeting N-terminal signal peptide. We focused on L2:DDRGK1 as it was enriched postnatally, neuron-specific, not annotated in ENSEMBL and experimentally validated by our 293T-based CRISPRa experiment (Fig. 4A; Fig. 5A; Supplemental Fig. S6B; Supplemental Table 9). GWAS studies have also identified a DDRGK1 associated risk locus for Parkinson’s disease (Nalls et al. 2014; Chang et al. 2017). The canonical DDRGK1 protein product is anchored to the ER membrane by an N-terminal 27 amino acid signal peptide (Fig. 5B) and plays a role in ER homeostasis and ER-phagy (Liang et al. 2020; Liu et al. 2017). In the predicted translated product of the L2:DDRGK1 TcGT, the signal peptide is replaced by a 10 amino acid L2-encoded sequence, conserved in new-world primates, but harboring non-synonymous substitutions in old-world primates (Fig. 5B; Supplemental Fig. S7A). Of note, this L2 integrant is absent in mice (Supplemental Fig. S7A). Furthermore, the L2:DDRGK1 TcGT is detected in the Rhesus Macaque developing brain with the same prenatal to postnatal expression dynamics as in humans (Supplemental Fig. S7B & C). We therefore transfected HEK293T cells with plasmids expressing HA-tagged versions of either the canonical “wild-type” (WT) DDRGK1 transcript or its TcGT counterpart and examined the subcellular localization of the resulting proteins by indirect immunofluorescence (Fig. 5C) and by cellular fractionation followed by western blotting (Fig. 5D). Confocal microscopy revealed that WT:DDRGK1-HA largely co-localized with BIP, an ER membrane marker, while L2:DDRGK1-HA displayed a diffuse cytosolic pattern (Fig. 5C). Cellular fractionation further confirmed that the WT DDRGK1 isoform was sequestered in the membrane fraction, whereas the L2:DDRGK1 counterpart was enriched in cytosol (Fig. 5D).

As N-truncated isoforms made up the largest category of *in-silico* predicted TcGT products besides full-length proteins, we next asked how widespread this type of TE-induced protein re-localisation might be. For this, we intersected a database of signal peptide-containing proteins with our initial list of 480 TcGT-encoded protein products (Fig. 5E; Supplemental Table 9). Of 94 TcGT products predicted to be N-truncated, 12 contained a putative signal peptide in the canonical isoform. This prediction was supported in 11 cases *in silico* by signalP 5.0 (Almagro Armenteros et al., 2019), which predicted that in all of these instances the TcGT isoforms lacked this putative signal peptide (Supplemental Fig. S8). Therefore, subcellular re-targeting may be a frequent consequence of TE-driven protein innovation.

## Discussion

An increasing number of studies are aimed at unravelling the transcriptional dynamics of human neurogenesis (Li et al. 2018; Cardoso-Moreira et al. 2019; Keil et al. 2018), yet, so far, little attention has been paid to the participation of TEeRS in this process. While retrotransposition of L1HS elements has been suggested to contribute to neuronal plasticity, experimental support for this model is lacking, and the vast majority of TEs hosted by the human genome have long lost the ability to spread (Muotri et al. 2005; Brouha et al. 2003). This prompted us to hypothesise that TEs might exert far greater influences on brain development through their ability to shape gene expression. As a first step towards testing this model, we analysed two independent human neurogenesis RNA-seq datasets with a ‘TE centric’ approach. This led us to uncover that the transposcriptome undergoes profound changes at each stage of brain development, with the expression of individual TE subfamilies largely anti-correlating to that of their cognate KZFP controllers. Strikingly, KZFP genes were globally downregulated at postnatal versus prenatal stages, coincident with the upregulation of their TE targets. Recent indications from an analysis of TEs resistant to loss of DNA methylation during the wave of epigenetic reprogramming in human primordial germ cells (hPGCs) showed modest anti-correlations of KZFPs and their target TE subfamilies in prenatal neurogenesis (Dietmann et al. 2020). The proposal that KZFPs may mediate the exaptation of TEs as developmental enhancers marked in hPGCs is intriguing and, combined with our analyses, suggests a multifaceted KZFP and TE mediated spatiotemporal transcriptional network, not only in prenatal stages but also highly prevalent after birth, with TEeRS playing important roles as alternative promoters, in addition to enhancers, throughout. Indeed, correlative expression studies on genic KZFP targets suggest that KZFPs may also directly regulate gene promoters during human neurogenesis independently from their TE binding ability (Farmiloe et al. 2020), and KZFPs were amongst genes previously found to be most differentially expressed between the chimpanzee and human brain (Nowick et al. 2009). Increasing evidence also supports a regulatory role for KZFP-targeted TEs in this and other developmental contexts (Ecco et al. 2016, 2017; Chen et al. 2019; Pontis et al. 2019; Turelli et al. 2020). For example, we recently demonstrated that two primate-restricted KZFPs, ZNF417 and ZNF587, control the expression of neuronal genes such as *PRODH* and *AADAT* via the regulation of HERVK-based TEeRS (Turelli et al. 2020). Furthermore, studies on the transcriptional co-repressors KAP1 and DNMT1 in hNPCs have highlighted their roles in the regulation of TEs and secondarily of cellular genes (Brattås et al. 2017; Jönsson et al. 2019). However, *in vitro* models do not recapitulate the global spatiotemporal complexity of gene and TE expression in the brain, nor its diverse cell type milieu throughout development, hence the interest of performing large scale ‘TE centric’ bioinformatics analyses on large post-mortem brain RNA-seq datasets.

De-repression of TEs, specifically of the LTR class, has been associated with various neurological disorders such as amyotrophic lateral sclerosis (ALS), Alzheimer’s disease (AD) and multiple sclerosis (MS) (Tam et al. 2019; Jönsson et al. 2020). The upregulation of LTR class elements in adult versus early prenatal brain is intriguing, as it suggests that LTR transposcription *per se* is a developmentally regulated feature of neurogenesis, which when deregulated is associated with a disease state. We propose that increased postnatal TE expression may possibly be reflective of the development of cell types not present in early prenatal stages, such as astrocytes, microglia and oligodendrocytes, the developmental and transcriptional trajectories of which were identified by scRNA-seq analyses (Li et al. 2018). To determine the transposcriptome in scRNA-seq data remains technically challenging because many TE-derived transcripts are lowly abundant, a limitation that will hopefully be alleviated by progress in sequencing techniques and computational approaches (Linker et al. 2020). Of note, TEs heavily contribute to long non-coding RNAs (lncRNAs), which are abundant in the human brain (Derrien et al. 2012; Kelley and Rinn 2012; Zimmer-Bensch 2019). It is plausible that upregulated TE transcripts play a role in this context, thereby exerting not cis-but trans-acting influences, the identification of which is far more challenging.

One increasingly well-characterised aspect of TE co-option is the engagement of TEeRS as alternative promoters. A wide range of oncogene-encoding TE-driven TcGTs have been documented in recent surveys of cancer databases (Jang et al. 2019; Attig et al. 2019), but the role of these transcript variants in physiological conditions remains largely undefined. Tissue-specific TcGTs have also been detected in the mouse developing intestine, liver, lung, stomach and kidney (Miao et al. 2020). Here, we demonstrate not only the spatially and temporally orchestrated expression of TcGTs in the developing human brain, but also that these TcGTs are largely organ- and cell type-specific. Some of them appear to be solely responsible for the expression of the involved gene, whereas others were present alongside canonical non-TE-driven transcripts, indicating sophisticated levels of regulation.

By experimental activation of a selected subset of antisense L2-driven TcGTs with CRISPRa and functional analyses of the product of the L2:DDRGK1 transcript, we highlight the functional relevance of this phenomenon for human neurogenesis. DDRGK1 is an ER membrane-associated protein with critical roles in UFMylation, an ubiquitin-like modification, and is involved in the unfolded protein response and ER-phagy (Liu et al. 2017; Liang et al. 2020). DDRGK1 is essential to target interactors like UFL1, the UFMylation ligase, to the ER membrane. The novel cytosolic chimeric L2:DDRGK1 protein, where a short N-terminal sequence derived from the L2 integrant replaces the signal peptide characteristic of its canonical counterpart, may therefore exert novel functions in the cytosol of postnatal to adult neurons. As signal peptide excision seems to affect a number of other TcGT products, this example may illustrate a more general phenomenon, whereby TE-driven genome evolution generates novel protein isoforms altering critical cell functions.

Our study indicates that the exaptation of TE-embedded regulatory sequences and its facilitation by TE-targeting KZFP controllers have significantly contributed to the complexity of transcriptional networks in the developing human brain. This warrants efforts aimed at delineating the evolutionary and functional impact of this phenomenon, and at defining how its alterations, notably in the context of inter-individual differences at these genomic loci, translates into variations in brain development, function and disease susceptibility.

## Methods

### Datasets

Raw RNA-seq fastq files for human and Rhesus macaque brain development (Cardoso-Moreira et al. 2019) were downloaded from the European Nucleotide Archive (datasets PRJEB26969 and PRJEB26956, respectively).

Raw RNA-seq fastq files for the GTEX and Brainspan (phs000424.v7.p2, phs000755.v2.p1), were downloaded from the dbGaP authorized access platform. Processed bed files containing regional neuronal or non-neuronal ATAC-seq peak loci from the Brain Open Chromatin Atlas (Fullard et al. 2018) were downloaded for hg19. To generate consensus neuronal and non-neuronal ATAC-peak bed files, bed coordinates from all regions were combined and overlapping peak coordinates merged using bedtools merge. Processed bed files for CAGE-seq peak loci from FANTOM5 (Forrest et al. 2014) were downloaded for hg19 (Lizio et al. 2015). Signal peptide containing proteins in human were downloaded from http://signalpeptide.com/index.php. Processed bed files from KZFP ChIP-exo experiments were used from our previous study (Imbeault et al. 2017).

### RNA-seq analysis

Reads were mapped to the human (hg19), or macaque (rheMac8) genome using hisat2 (Kim et al. 2015) with parameters hisat2 -k 5 --seed 42. Counts on genes and TEs were generated using featureCounts (Liao et al. 2014). To avoid read assignation ambiguity between genes and TEs, a gtf file containing both was provided to featureCounts. For repetitive sequences, an in-house curated version of the Repbase database was used (fragmented LTR and internal segments belonging to a single integrant were merged), generated as previously described (Turelli et al. 2020). Minor modifications to the repeat merging pipeline described in Turelli et al., 2020 were made for Macaque (RepeatMasker 4.0.5 20160202) with the distance between two LTR elements of the same orientation to an ERV-int fragment being less than 400bp. For genes the ensemble release 75 annotation was used. Only uniquely mapped reads were used for counting on genes and TEs with the command ‘featureCounts - t exon -g gene_id -Q 10’. For the Brainspan dataset, samples with less than 10 million unique mapped reads on genes were discarded from the analysis. TEs that did not have at least one sample with 50 reads or overlapped an exon were discarded from the mapping TE integrant analysis. For estimating TE subfamilies expression level, reads were summarized using the command featureCounts -M -- fraction -t exon -g gene_id -Q 0 then, for each subfamily, counts on all TE members were added up. As the Cardoso-Moreira et al., 2019 RNA-seq was stranded data, reads on both strands were combined for TEs to facilitate comparison to the non-stranded Brainspan dataset. Normalization for sequencing depth was done for both genes and TEs using the TMM method as implemented in the limma package of Bioconductor (Gentleman et al. 2004) and using the counts on genes as library size. Differential gene expression analysis was performed using voom (Law et al. 2014) as it has been implemented in the limma package of Bioconductor (Gentleman et al. 2004). A gene (or TE) was considered to be differentially expressed when the fold change between groups was greater than two and the p-value was smaller than 0.05. A moderated t-test (as implemented in the limma package of R) was used to test significance. P-values were corrected for multiple testing using the Benjamini-Hochberg’s method (Benjamini and Hochberg 1995). Temporal expression correlation analyses of individual genes, TE integrants or subfamilies were performed between Brainspan and Cardoso datasets using the ‘Pearson’ method. For inter-regional correlations within the Brainspan dataset, only expressed genes or TEs common to all regions were considered. Bam files and sashimi plots were visualised using the Integrative Genomics Viewer (Katz et al. 2015; Robinson et al. 2011).

### TcGT detection pipeline

First, a per sample transcriptome was computed from the RNA-seq bam file using Stringtie (Kovaka et al. 2019) with parameters -j 1 -c 1. Each transcriptome was then crossed using BEDTools (Quinlan and Hall 2010), to ensembl hg19 (or rheMac8) coding exons and curated RepeatMasker to extract TcGTs with one or more reads spliced between a TE and genic exon for each sample. Second, a custom python program was used to annotate and aggregate the sample level TcGTs into counts per stages (defined in Supplemental Fig. S1B). In brief, for each dataset, a GTF containing all annotated TcGTs was created and TcGTs having their first exon overlapping an annotated gene, or TSS not overlapping a TE were discarded. From this filtered file, TcGTs associated with the same gene and having a TSS within 100bp of each other were aggregated. Finally, for each aggregate, its occurrence per group was computed and a consensus transcript was generated for each TSS aggregate. For each exon of TcGT aggregate, its percentage of occurrence across the different samples was computed and integrated in the consensus if present in more than 30% of the samples the TcGT was detected in. All samples available in both datasets were used regardless of mapped read count.

From the resulting master file, additional criteria were applied to determine prenatal, postnatal or continually expressed TcGTs. 1. Only TcGTs that were present in at least 20% of prenatal, postnatal or 20% of both pre and postnatal samples (continual) were kept for each dataset. 2. To ensure TcGTs were robustly detectable in the different datasets, TcGT files were merged based on the same TSS TE and associated gene name. 3. TcGTs were required to exhibit the same temporal transcriptional behaviour in both datasets. I.E a 2 fold change in TcGT detection pre vs postnatal and vice versa or a lower fold change in both datasets (continual). This resulted in the 480 robustly detectable temporal TcGTs in Fig. 3A and Supplemental Table 9. These TcGTs were further filtered for strong promoter regions using a Bedtools intersect of the 200bp up and downstream of the TcGT TSS with FANTOM5 CAGE-seq (Forrest et al. 2014) and BOCA neuronal and non-neuronal consensus ATAC-seq peak bed files (Fullard et al. 2018). TcGT TSS loci were also intersected with ENSEMBL (GRCh37.p13) transcriptional start sites to determine non-annotated transcripts.

### Protein product prediction

DNA sequences were retrieved for each TcGTs consensus and protein products were derived from the longest ORF in the three reading frames using biopython (Cock et al. 2009). The resulting translation products were aligned against the protein sequence of the most similar cognate gene isoforms (exons intersect between TcGTs and each gene isoform) and classified into several categories. Proteins with no alignment for any isoform were classified as out-of-frame, therefore not clear or not aligned. In-frame peptides were further classified according to their N-terminal modifications: Normal, TcGT ORF peptides align perfectly with cognate ORF peptides; N-add, TcGT ORF peptides encode novel in-frame N-terminal amino acids followed by the full length cognate protein sequence; N-truncated, TcGT ORF peptides lack parts of the cognate N-terminal protein sequence and might contain novel in-frame N-terminal amino acids. TcGTs that we could not clearly classify were grouped in the ‘other’ category, such as TcGTs including C-terminal modifications. If the classification was ambiguous for different protein isoforms, the normal category was always privileged.

### TE and KZFP age estimation

TE subfamily ages were downloaded from DFAM (Hubley et al. 2016). To compare KZFP ages we developed a score we called Complete Alignment of Zinc Finger (CAZF) (as we described in Thorball et al. 2020), which rely on the alignments of zinc finger domains, using only the four amino acid presumably touching DNA. Briefly, alignment scores made with BLOSUM80 matrix were used, normalised by the ‘perfect’ alignment score (alignment against itself) and by the length of the alignment. To compute an age for KZFPs, we relied on inter-species clusters of KZFPs made with CAZF score. KZFPs with CAZF>0.5 were clustered together, using a bottom-up approach. The divergence time between human and the farthest species present in the cluster was used as the age of individual KZFPs in the cluster. Multiz alignments for L2:DDRGK1 locus were extracted from the UCSC genome browser

### Cell culture

Human embryonic kidney 293T (HEK293T) cells and SH-SY-5Y neuroblastoma cells were cultured in DMEM supplemented with 10% fetal calf serum and 1% penicillin/streptomycin.

### Transfection

Transient transfection of HEK293T cells was performed with FuGENE HD (Promega) as per the manufacturer’s recommendation. Cells were harvested 48 hours after transfection for either RNA extraction or immunofluorescence.

### CRISPRa

The SP-dCas9-VPR (Addgene 63798) (Chavez et al. 2015) and the gRNA cloning vector (Addgene 41824) (Mali et al. 2013) were gifts from George Church. gRNAs were designed with CRISPOR (Concordet and Haeussler 2018), using input DNA sequence 50 to 300bp upstream of the TE resident CAGE peak and the most 5’ location of RNA-seq reads mapping to the TcGT TE TSS loci. Multiple gRNAs were selected for each TcGT to control for gRNA specific effects and increase experimental robustness. gRNA oligonucleotides were synthesised (Microsynth) with the recommended overhangs (Supplemental Table 10) for integration into the gRNA cloning vector (Mali et al. 2013). gRNA oligonucleotides were annealed and extended using Phusion High Fidelity DNA polymerase master mix (NEB) with thermal cycling conditions of 98°C two minutes (1x), 98°C 10 seconds + 72°C 20 seconds (3x) and 72°C for five minutes. 10μg of SP-dCas9-VPR was digested with Af1II (NEB) in CutSmart buffer for two hours at 37°C, followed by gel electrophoresis and purification of the correct sized band of linearised plasmid with E.Z.N.A Gel Extraction Kit (Omega Bio-tek). The resulting linearised plasmid and double stranded oligonucleotides were ligated using Gibson Assembly Master Mix (NEB) as per manufacturer’s recommendations. The resulting gRNA containing plasmid was transformed into HB101 chemically competent *E.coli,* with colonies containing the transformed plasmid selected on agar plates containing kanamycin, followed by colony picking for growth in kanamycin agar broth followed by GeneJET Plasmid Miniprep (ThermoFisher). gRNA plasmids were Sanger sequenced to detect the correct insertion of specific gRNA sequences. 300,000 HEK293T cells were seeded per well of a six well plate. 24 hours later, co-transfection was performed with 1μg each of SP-dCas9-VPR and TcGT targeting gRNA containing gRNA cloning vector. SP-dCas9-VPR or empty gRNA cloning vector alone were transfected as non-targeting controls. Cells were harvested for RNA 48 hours post-transfection.

### RT-PCR and qRT-PCR

Primers to detect TcGTs were designed with Primer3 (Untergasser et al. 2012) by inputting DNA sequences covering and flanking the splice junction between the TE and genic exon (Supplemental Table 10). One primer was required to be present in the TE sequence where RNA-seq reads were detected downstream of a CAGE-peak, whilst the other was present in the first or second genic exon. BLAT (Kent 2002) of primer sequences against the human genome ensured only uniquely mapping primers were used. RNA was extracted from cells using the NucleoSpin RNA mini kit (Macherey-Nagel) with on-column deoxyribonuclease treatment. 1ug RNA was used in the cDNA synthesis reaction with the Maxima H minus cDNA synthesis master mix (ThermoFisher) and RT-PCR was performed with Phusion High Fidelity DNA polymerase master mix (NEB) each with the manufacturer recommended PCR thermal cycles, on a 9800 Fast Thermal Cycler (Applied Bioscience). PCR products were visualised by 1.5% agarose gel electrophoresis stained with SYBR Safe DNA gel stain (ThermoFisher) and imaged with a BioDoc-It imaging system (UVP). Bands of the correct size were excised, gel purified with E.Z.N.A Gel Extraction Kit (Omega Bio-tek) and Sanger sequenced using primers used for PCR. The correct PCR product was confirmed using BLAT (Kent 2002) of the Sanger sequencing results against the human genome. qRT-PCR was performed with PowerUp SYBR Green Master Mix on a QuantStudio 6 Flex Real-Time PCR system. The standard curve method was used to quantify expression normalised to *BETA ACTIN* with no amplification in the no reverse transcriptase control.

### Cloning of WT:DDRGK1 and L2:DDRGK1

The WT:DDRGK1 cDNA Clone (Genbank accession:HQ448262 ImageID:100071664) was obtained from the ORFeome Collaboration (http://www.orfeomecollaboration.org/) in the *pENTR223* vector without a stop codon. The L2:DDRGK1 sequence was PCR amplified with Phusion High Fidelity DNA polymerase master mix (NEB), using cDNA generated in the L2:DDRGK1 CRISPRa experiment with gRNA 1. This ensured the *bona fide* L2 driven transcript was cloned. Cloning primers used are shown in Supplemental Table 10, with the forward primer containing a CACC Kozak sequence and the reverse primer omitting the stop codon. Thermal cycling conditions were 98°C 30 seconds (1x), 98°C 10 seconds + 60°C 15 seconds + 72°C 15 seconds (35x) and 72°C for 10 minutes. A 466bp PCR fragment was extracted after agarose gel electrophoresis, purified with E.Z.N.A Gel Extraction Kit (Omega Bio-tek), transformed into chemically competent HB101 E.coli, colonies picked and mini-prepped. WT:DDRGK1 and L2:DDRGK1 in the *pENTR* vectors were then shuttled into pTRE-3HA (Imbeault et al. 2017) with the Gateway LR Clonase II Enzyme mix (ThermoFisher) as per manufacturer’s instructions. pTRE-3HA produces proteins with three C-terminal HA tags in a doxycyclin-dependent manner.

### Cellular fractionation

Approximately 400,000 HEK293T cells in different wells of a 6 well plate were transfected with either pTRE-WT:DDRGK1-HA or pTRE-L2:DDRGK1-HA whose expression was induced for 48 hours by adding 1μg/ml doxycycline to the media. After 48 hours wells were washed with 1ml ice cold PBS and cells were scraped and transferred to Eppendorf tubes on the second wash. After centrifugation at 300rcf for five minutes at 4°C, PBS was aspirated, cells re-suspended in 400μl ice-cold cytoplasmic isolation buffer (10mM KOAc, 2mM MgOAC, 20mM HEPES pH7.5, 0.5mM DTT, 0.015% digitonin) and centrifuged at 900rcf for five minutes at 4°C. Supernatant was collected as the cytoplasmic fraction and the remaining pellet was re-suspended in 400μl of membrane isolation buffer (10mM HEPES, 10mM KCl, 0.1mM EDTA pH8, 1mM DTT, 0.5% Triton X-100, 100mM NaF), then centrifuged for 10 minutes at 900rcf at 4°C to pellet nuclei with the supernatant collected as the membrane fraction. Pelleted nuclei were resuspended in 400μl of lysis buffer (1% NP-40, 500mM Tris-HCL pH8, 0.05% SDS, 20mM EDTA, 10mM NaF, 20mM benzamidine) for 10 minutes on ice, centrifuged for 10 minutes at 900rcf at 4°C and the supernatant collected as the nuclear fraction. 100μl of 4x NuPAGE LDS sample buffer (ThermoFisher) was added to the 400μl cellular fractions and samples boiled at 95°C for five minutes.

### Western blot

20μl of each cellular fraction was used for SDS-PAGE in a NuPAGE 4-12% Bis-TRIS gel and MOPs running buffer (ThermoFisher). For subcellular fraction marker proteins, the same amount of lysate was added from each sample but for the HA blot, pTRE-WT:DDRGK1-HA samples were diluted 1:50 due to high over-expression levels compared to pTRE-L2:DDRGK1-HA. Proteins were transferred to a nitrocellulose membrane using an iBLOT 2 dry blotting system (ThermoFisher) and analysed by immunoblotting using CALNEXIN (Bethyl A303-696A, 1:2000), LAMIN B1 (Abcam ab16048, 1:1000), β TUBULIN (Sigma T4026, 1:1000), HA-HRP conjugated (Roche 12013819001, 1:2000). HRP-conjugated anti-mouse (GE Healthcare NA931V, 1:10000) and HRP-conjugated anti-rabbit (Santa Cruz sc-2004 1:5000) antibodies were used where appropriate and the blot was visualised using the Fusion SOLO S (Vilber).

### Immunofluorescence

HEK293T cells were plated on glass coverslips and immunofluorescence was performed as previously described (Helleboid et al. 2019) 48 hours post-transfection and expression induction with 1μg/ml doxycycline for pTRE-WT:DDRGK1-HA or pTRE-L2:DDRGK1-HA. Once 70% confluent, cells were washed three times with PBS, fixed in ice-cold methanol for 20 minutes at −20°C then washed three more times with PBS. Cells were blocked with 1% BSA/PBS for 30 minutes and then incubated with antibodies for HA.11 (BioLegend MMS-101P, 1:2000) and BIP (Abcam ab21685, 1:1000) in 1% BSA/PBS for one hour. Three washes with PBS were performed, followed by incubation with anti-mouse and anti-rabbit Alexa 488 or 568 (ThermoFisher 1:800) for one hour. DAPI (1:10000) was added in the last 10 minutes of incubation, samples washed three times with PBS and coverslips mounted on slides with ProLong Gold Antifade Mountant (ThermoFisher). Images were acquired on a SP8 upright confocal microscope (Leica) and processed in ImageJ.

### Data Access

No additional high throughput data was generated in this study.

## Supporting information

Playfoot_Supplemental_Figs

## Acknowledgements

We thank all members of the Trono Lab for helpful and insightful discussions, along with Samuel Corless and Nezha Benabdallah for critical reading of the manuscript.

## Author Contributions

C.P. and D.T. conceived the study, interpreted the data, and wrote the manuscript. C.P. performed bioinformatics analyses and all experiments. J.D. and S.S. developed key code and performed bioinformatics analyses. S.D. performed the GTEX TcGT analysis and *in silico* translation of TcGTs. A.C. performed the KZFP aging analysis and determined KZFP TE subfamily targets. E.P. contributed to bioinformatics tools and code. All authors reviewed the manuscript.

