## Supplementary material for "Transposable elements and their KZFP controllers are drivers of transcriptional innovation in the developing human brain": Playfoot_Supplemental_Figs

Playfoot_Fig. S1

A

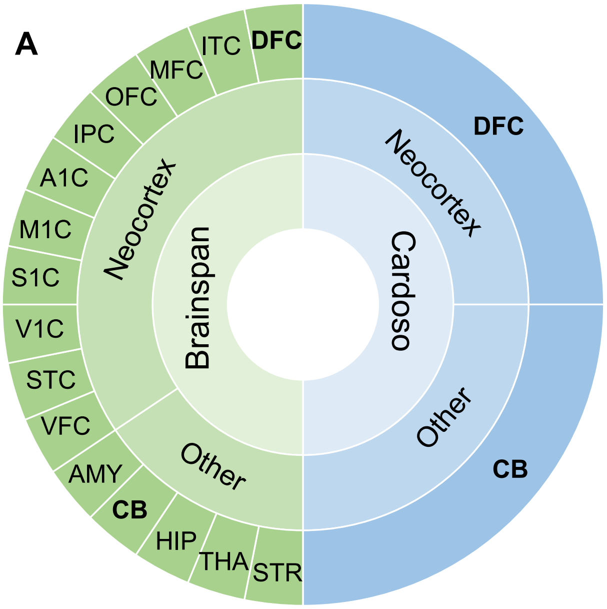


B

| Brainspan Region | Abv. | 1 (4-7pcw) | 2A (8-9pcw) | 2B (10-12pcw) | 3A (13-15pcw) | 3B (16-18pcw) | 4 (19-24pcw) | 5 (25-38pcw) | 6 (Birth-5month) | 7 (6-18 month) | 8 (19month-5yr) | 9 (6-11 yr) | 10 (12-19 yr) | 11 (20-60+ yr) | Sum |
|---|---|---|---|---|---|---|---|---|---|---|---|---|---|---|---|
| Amygdala | AMY | 0 | 2 | 4 | 4 | 1 | 2 | 1 | 3 | 1 | 1 | 2 | 3 | 7 | 31 |
| Cerebellar cortex | CB | 0 | 0 | 3 | 2 | 1 | 3 | 2 | 2 | 2 | 3 | 2 | 4 | 7 | 31 |
| Dorsolateral prefrontal cortex | DFC | 0 | 2 | 4 | 4 | 2 | 4 | 1 | 2 | 2 | 2 | 2 | 3 | 6 | 34 |
| Hippocampus | HIP | 0 | 2 | 4 | 4 | 2 | 4 | 1 | 3 | 1 | 1 | 2 | 2 | 7 | 33 |
| Inferior temporal cortex | ITC | 0 | 1 | 4 | 4 | 0 | 3 | 1 | 3 | 2 | 2 | 2 | 4 | 7 | 33 |
| Medial prefrontal cortex | MFC | 0 | 2 | 3 | 4 | 2 | 4 | 1 | 2 | 2 | 2 | 2 | 3 | 6 | 33 |
| Mediodorsal nucleus of the thalamus | THA | 0 | 0 | 4 | 2 | 2 | 4 | 1 | 3 | 2 | 2 | 1 | 2 | 7 | 30 |
| Orbital prefrontal cortex | OFC | 0 | 2 | 4 | 3 | 1 | 3 | 1 | 2 | 2 | 1 | 2 | 3 | 7 | 31 |
| Posterior inferior parietal cortex | IPC | 0 | 0 | 4 | 4 | 2 | 3 | 1 | 2 | 3 | 2 | 2 | 4 | 7 | 34 |
| Primary auditory (A1) cortex | A1C | 0 | 0 | 4 | 4 | 2 | 2 | 1 | 3 | 1 | 2 | 2 | 2 | 7 | 30 |
| Primary motor (M1) cortex | M1C | 0 | 0 | 4 | 3 | 0 | 2 | 1 | 3 | 1 | 2 | 1 | 3 | 7 | 27 |
| Primary somatosensory (S1) cortex | S1C | 0 | 0 | 4 | 3 | 0 | 2 | 1 | 3 | 3 | 2 | 1 | 4 | 7 | 30 |
| Primary visual (V1) cortex | V1C | 0 | 0 | 4 | 4 | 2 | 4 | 1 | 2 | 2 | 1 | 2 | 4 | 6 | 32 |
| Striatum | STR | 0 | 0 | 4 | 4 | 2 | 3 | 1 | 2 | 1 | 2 | 1 | 1 | 7 | 28 |
| Superior temporal cortex | STC | 0 | 1 | 3 | 3 | 2 | 4 | 1 | 3 | 2 | 3 | 2 | 3 | 7 | 34 |
| Ventrolateral prefrontal cortex | VFC | 0 | 1 | 4 | 4 | 2 | 4 | 1 | 2 | 2 | 3 | 2 | 4 | 7 | 36 |
| | | | | | | | | | | | | | | | **507** |

| Cardoso Region | Abv. | 1 | 2A | 2B | 3A | 3B | 4 | 5 | 6 | 7 | 8 | 9 | 10 | 11 | Sum |
|---|---|---|---|---|---|---|---|---|---|---|---|---|---|---|---|
| Dorsolateral prefrontal cortex | DFC | 8 | 7 | 6 | 3 | 3 | 5 | 0 | 3 | 2 | 3 | 2 | 4 | 9 | 55 |
| Cerebellar cortex | CB | 12 | 6 | 9 | 3 | 3 | 0 | 0 | 5 | 3 | 2 | 3 | 4 | 9 | 59 |
| | | | | | | | | | | | | | | | **114** |

C

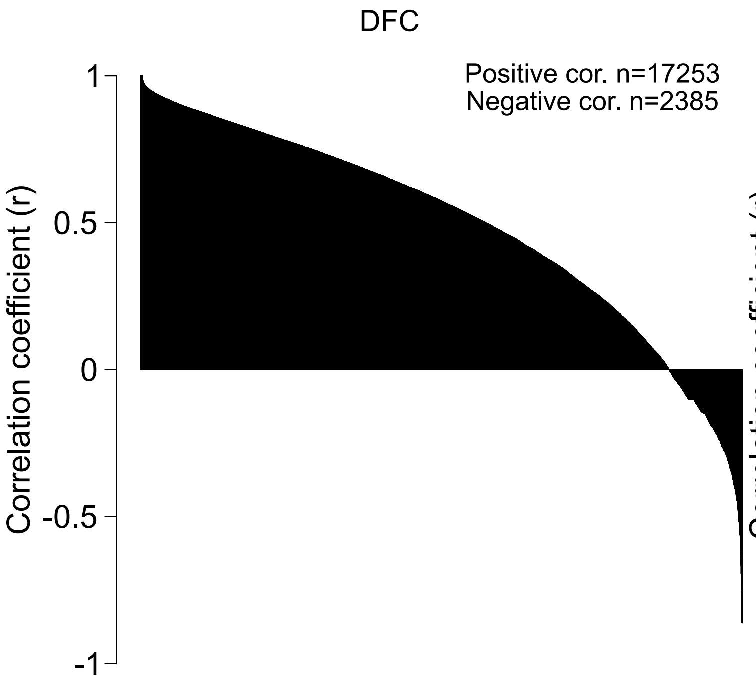


D

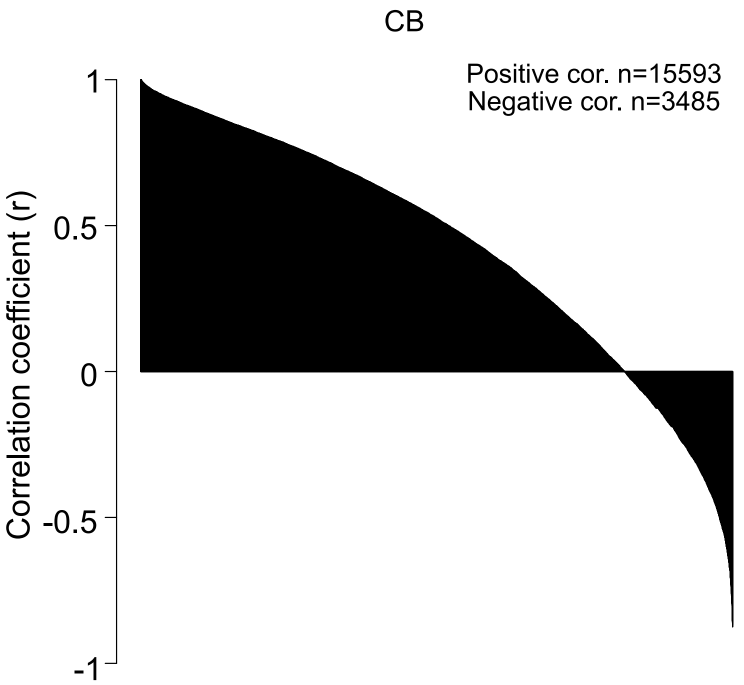

### Playfoot_Fig. S2

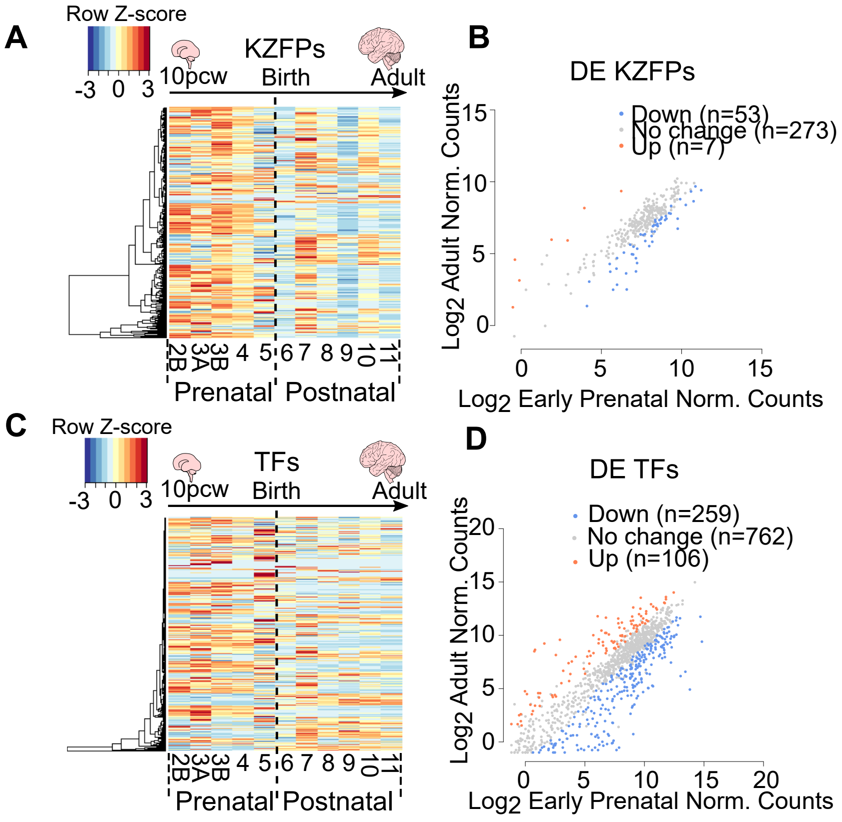

Playfoot_Fig. S3

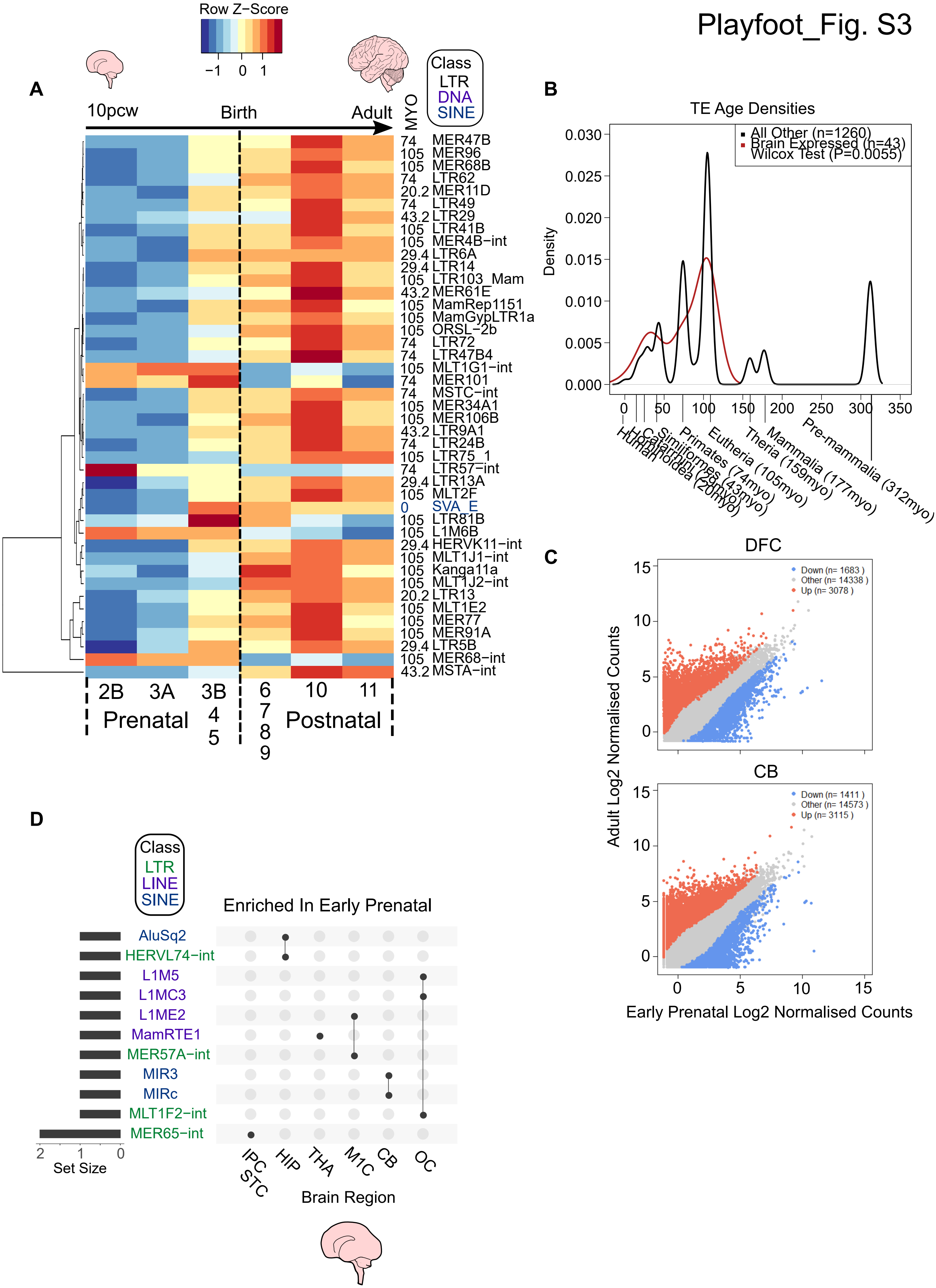

Playfoot_Fig. S4

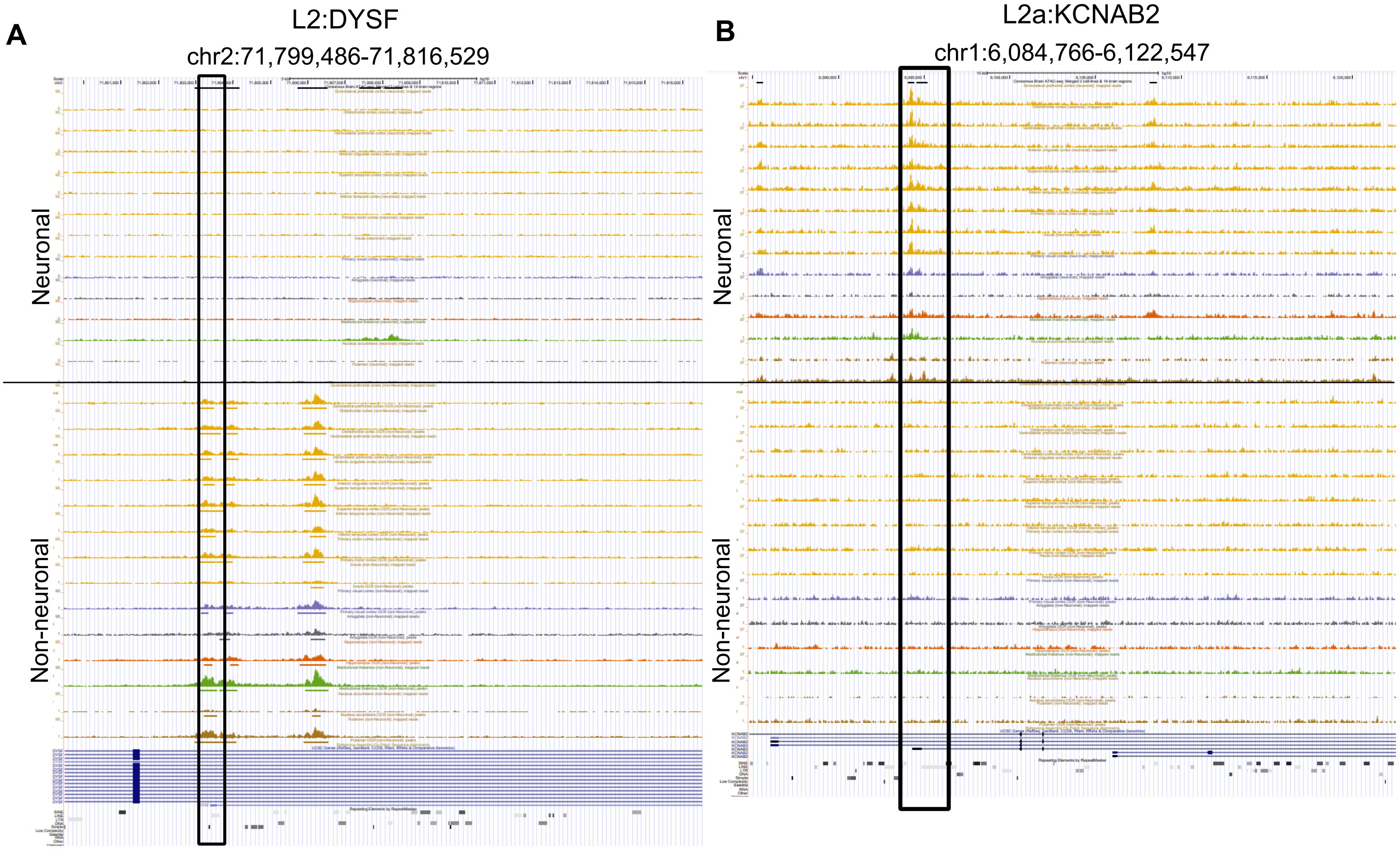

Playfoot_Fig. S5

A

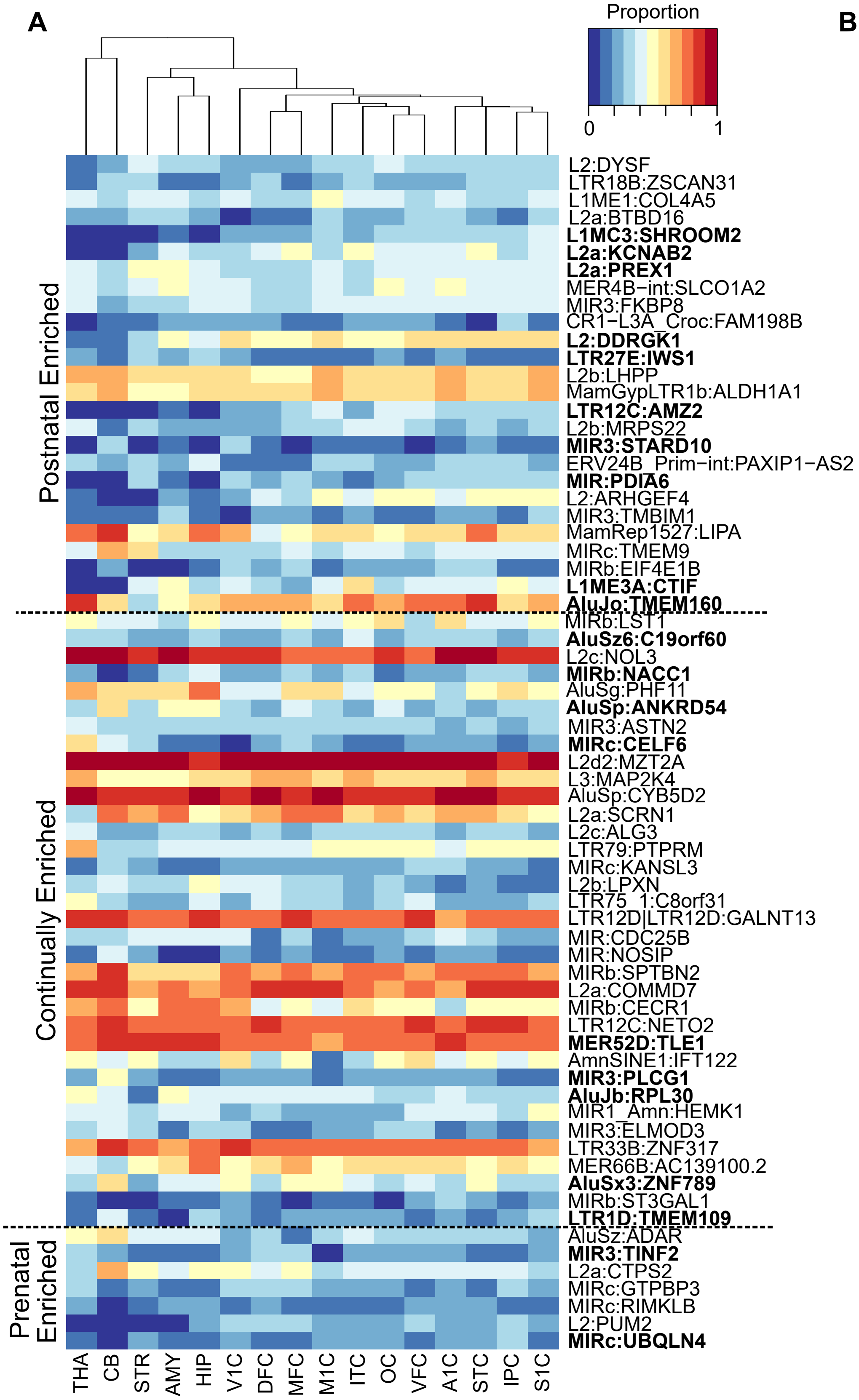


B

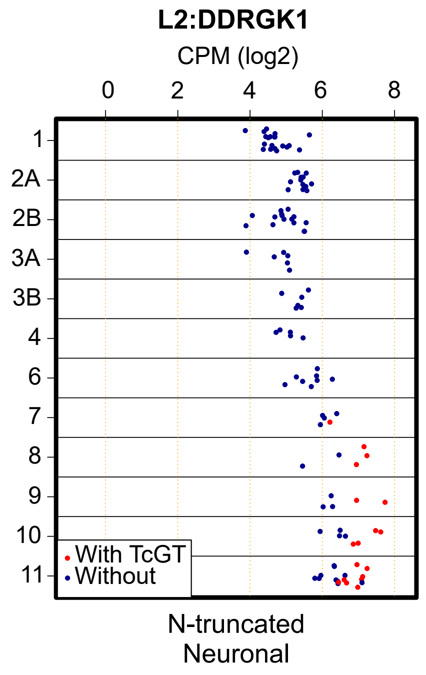

Playfoot_Fig. S6

A

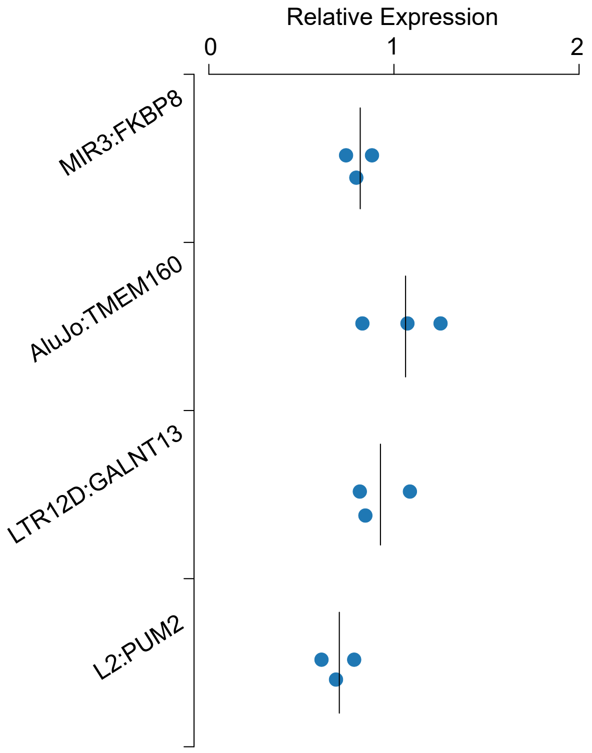


B

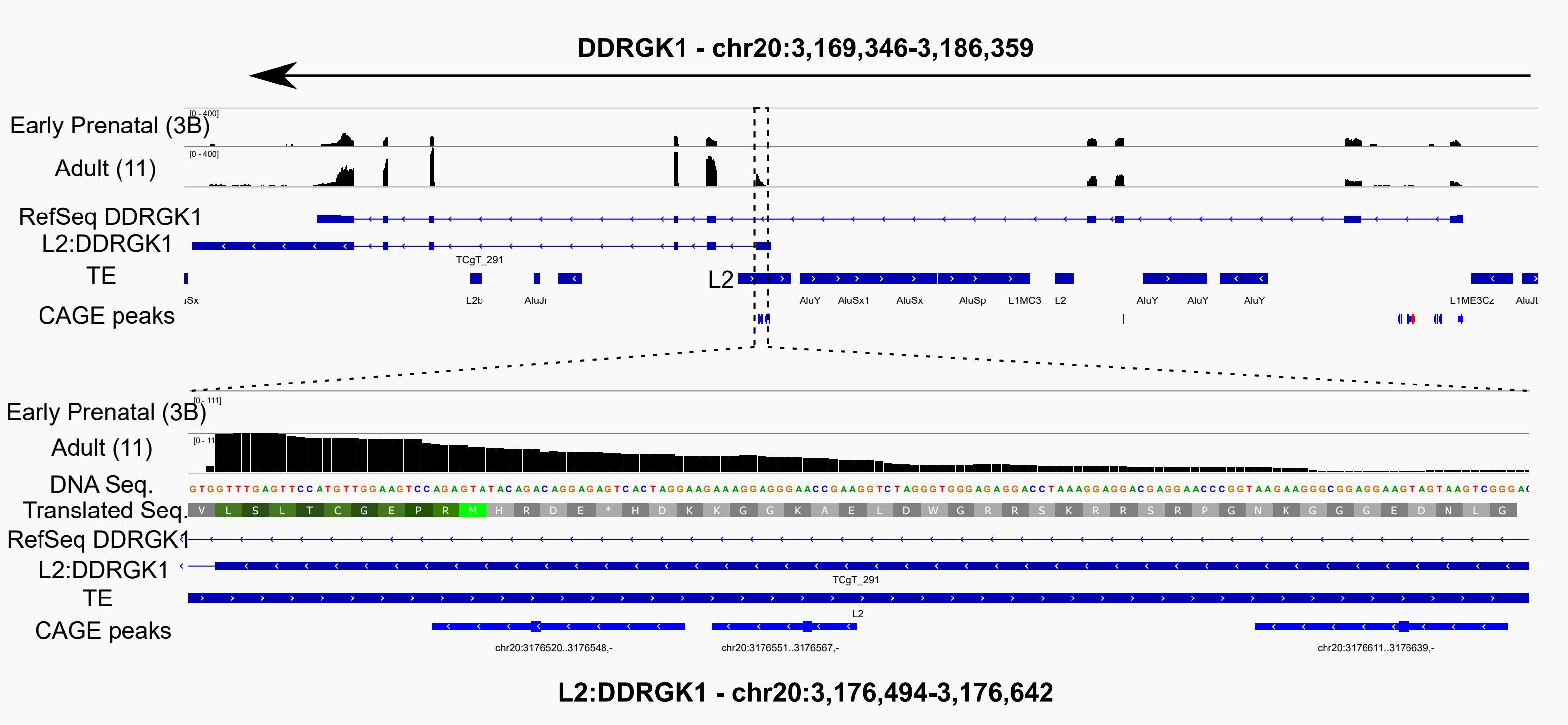

Playfoot_Fig. S7

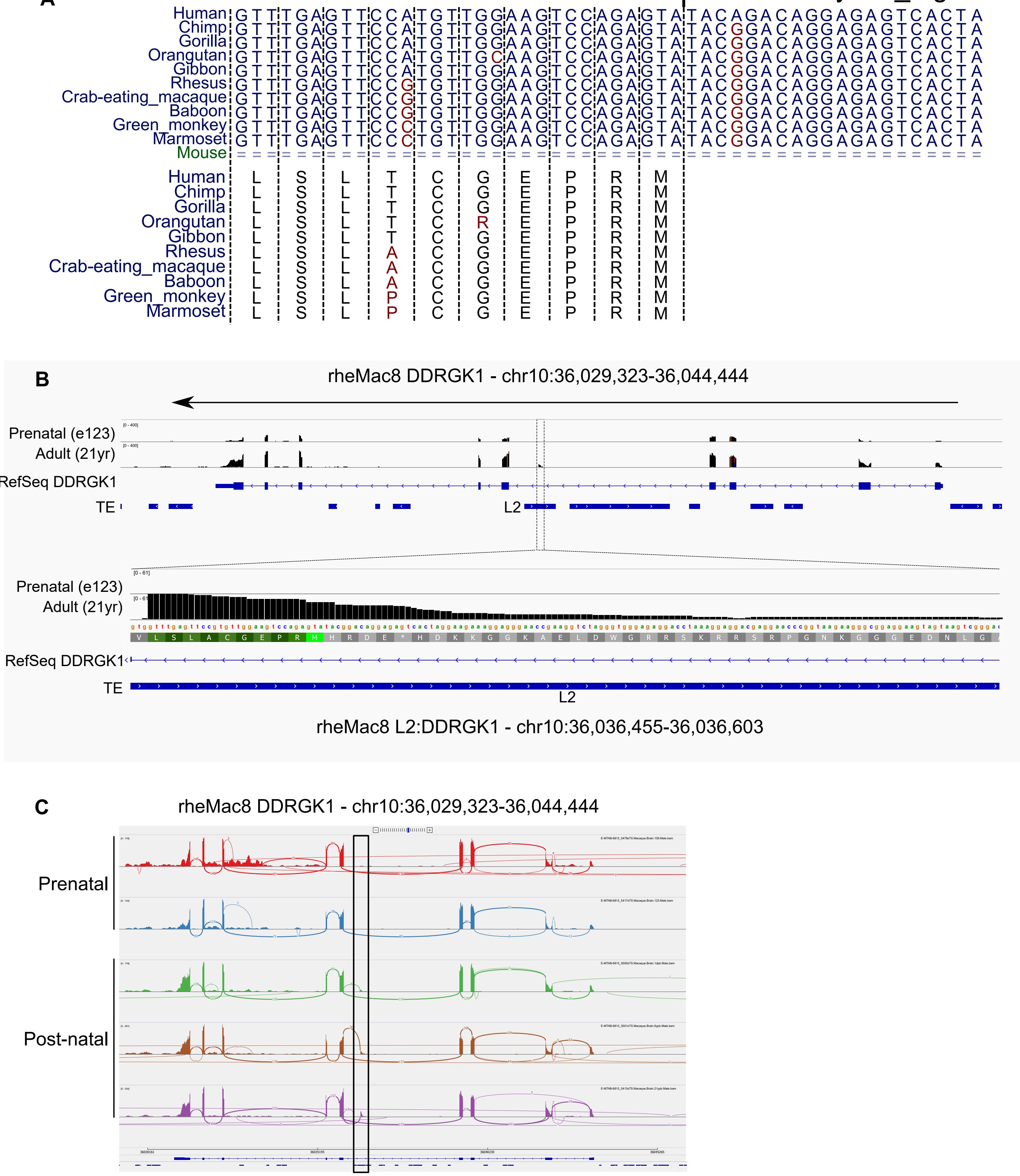

Playfoot_Fig. S8

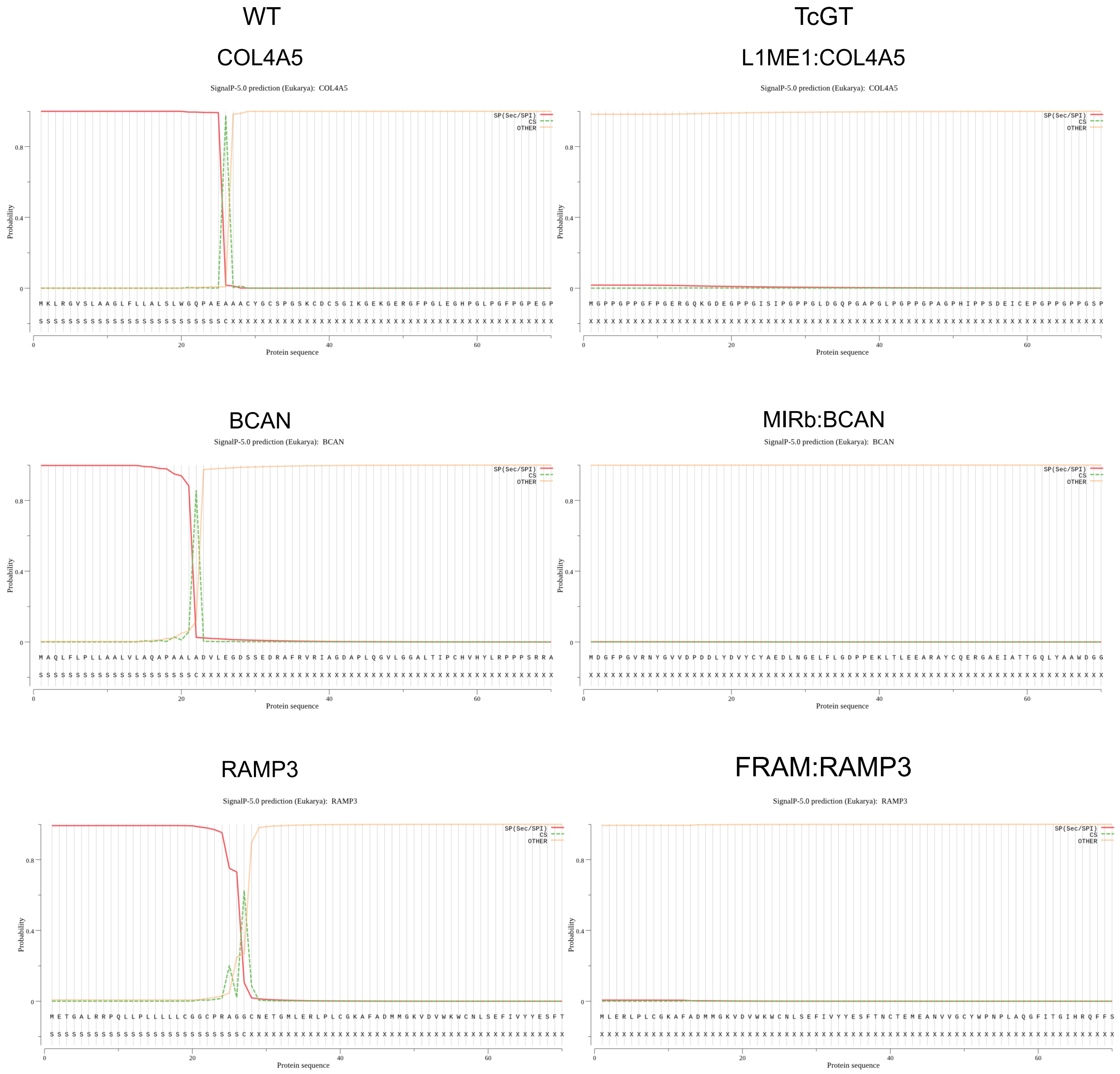

#### Supplemental Figure Legends:

**Figure S1. Summary of brain developmental datasets analysed.** (A) Donut plot of brain regions incorporated in both datasets. Region abbreviations are shown in B. (B) Number of samples analysed in both datasets, highlighting numbers of samples per region and stage. Stages were as defined by the Brainspan consortia. (C) Barplots depicting the Pearson correlation coefficient between Brainspan and Cardoso datasets for all genes passing expression criteria (methods) for the DFC and the CB (D).

**Figure S2. KZFP expression trajectory is different in the CB compared to the DFC.** (A) Heatplots of KZFP expression across human neurogenesis in the cerebellum. Scale represents the row Z-score. Stage 2A was omitted due to lack of samples for CB (see Fig. S1B). See also Supplemental Table 2. (B) Dot plot of differential expression analysis of KZFPs in the CB comparing adult (stage 11) to early prenatal stages (stage 2A to 3B) of neurogenesis. Only KZFPs behaving the same in both datasets are shown. Up (orange) represents KZFPs significantly upregulated in adult versus early prenatal (Fold change ≥ 2, FDR ≤ 0.05). Down (blue) represents KZFPs significantly downregulated in adult (Fold change ≤ -2, FDR ≤ 0.05). See also Supplemental Table 3. (C) Heatplots of TF expression across human neurogenesis in the CB. Scale represents the row Z-score. Stage 2A was omitted due to lack of samples for CB (see Fig. S1B). See also Supplemental Table 2. (D) Dot plot of differential expression analysis of TFs in the CB comparing adult (stage 11) to early prenatal stages (stage 2A to 3B) of neurogenesis. Only TFs behaving the same in both datasets are shown. Up (orange) represents TFs significantly upregulated in adult versus early prenatal (Fold change ≥ 2, FDR ≤ 0.05). Down (blue) represents TFs significantly downregulated in adult (Fold change ≤ -2, FDR ≤ 0.05). See also Supplemental Table 3. All plots show expression data from Brainspan.

**Figure S3. TE subfamilies and unique loci exhibit spatiotemporal expression patterns in the cerebellum.** (A) Heatplot of TE subfamilies with concordant expression behaviours between both datasets (Pearson correlation coefficient ≥ 0.7) across human neurogenesis in the cerebellum. See also Supplemental Table 4. The mean expression values for stages 3B, 4 and 5, and also stages 6, 7, 8 and 9 were combined and averaged to reduce inherent variability due to low numbers of samples for some stages (see Supplemental Fig. S1B). Scale represents the row Z-score. TE subfamily age in million years old (MYO) and class is shown to the right of the plot. (B) Density plot depicting estimated age of TEs in A ($P \leq 0.05$, Wilcoxon test). Evolutionary stages and corresponding ages are shown beneath the plot. (C) Dot plots of differential expression analysis of unique TE loci in the DFC and CB comparing adult (stage 11) to early prenatal stages (stage 2A to 3B) of neurogenesis. Only TEs behaving the same in both datasets are shown. Up (orange) represents TEs significantly upregulated in adult versus early prenatal (Fold change ≥ 2, FDR ≤ 0.05). Down (blue) represents TEs significantly downregulated in adult (Fold change ≤ -2, FDR ≤ 0.05). See also Supplemental Table 7 & 8. (D) UpSet plot showing the significantly enriched differentially expressed subfamilies for early pre-natal versus adult stages per region from unique mapping analyses. Set size represents the number of regions the specific TE was significantly differentially enriched in. Joined points represent combinations of significantly differentially expressed TE subfamilies. See also Supplemental Table 7 & 8.

**Figure S4. TcGTs are cell type specific.** (A) UCSC genome browser shots of TcGT TE TSS loci (black box) with consensus ATAC-seq peaks from isolated neuronal and non-neuronal cells from different regions of the adult human brain from BOCA (Fullard and Hauberg et al., 2018). Tracks are shown for the non-neuronal associated TcGT L2:DYSF and the neuronal associated TcGT L2a:KCNAB2. Track colours correspond to the following regions: Yellow = different neocortex regions, purple = primary visual cortex, black = amygdala, red = hippocampus, green = mediodorsal thalamus, brown= nucleus accumbens and putamen.

**Figure S5. TcGTs are spatially expressed in broad or specific brain regions.** (A) Heatplot showing the proportion of samples per brain region the 68 TcGTs were detected in the Brainspan dataset (same order as Fig. 4A). (B) Dot plots showing the gene expression level per stage for L2:DDRGK1 for samples where the TcGT was detected (red) and where it was not (blue) from Cardoso dataset as comparison to Fig. 4A.

**Figure S6. TcGTs are expressed in SH-SY-5Y neuroblastoma cells and L2:DDRGK1 is a predicted chimeric protein.** (A) qRT-PCR expression plots of the indicated TcGTs relative to *BETA ACTIN* using primers designed within the TcGT TE TSS and the first exon of the TcGT associated gene. (n=3 independent cultures of SH-SY-5Y cells). (B) Integrated genome viewer (IGV) image of the L2:DDRGK1 TcGT locus showing representative RNA-seq read pile-ups from early prenatal (stage 3B) and adult (stage 11). Zoom in highlights the DNA sequence and amino acid sequence with the L2 derived start codon highlighted in light green with subsequent peptides in dark green. Cage peaks are also shown. The gene and TcGT transcript is in the antisense strand orientation (right to left) whereas the L2 element is in the sense strand orientation (left to right).

**Figure S7. The L2:DDRGK1 TcGT is conserved in primates and has the same behaviour in macaque as in humans.** (A) Multiz DNA sequence alignments of the L2 element coding for the chimeric DDRGK1 TcGT isoform. The methionine start codon is indicated by the square arrow. Some of the L2 5' UTR was omitted for clarity but contained no overt sequence differences. The *in silico* translated product is shown below the DNA sequence, with dashed lines denoting codons. (B) Integrated genome viewer (IGV) image of the macaque L2:DDRGK1 TcGT locus showing representative RNA-seq read pile-ups from prenatal (embryonic day 123) and adult (21 years). Zoom in highlights the DNA sequence and translated amino acid sequence with the L2 derived start codon highlighted in light green with subsequent peptides in dark green. The gene and TcGT transcript is in the antisense strand orientation (right to left) whereas the L2 element is in the sense strand orientation (left to right).

**Figure S8. The signal peptide is lost in N-truncated TcGTs.** Plots generated from SignalP 5.0 showing computationally determined N-terminal signal peptide sequence and cleavage sites for canonical gene transcripts and for consensus N-truncated TcGT derived transcripts. Red line denotes the predicted signal peptide, dashed green line the predicted cleavage site and orange represents non-signal peptide sequence.
